# Structural and Functional Analyses Explain Pea KAI2 Receptor Diversity and Reveal Stereoselective Catalysis During Signal Perception

**DOI:** 10.1101/2021.01.06.425465

**Authors:** Angelica M. Guercio, Salar Torabi, David Cornu, Marion Dalmais, Abdelhafid Bendahmane, Christine Le Signor, Jean-Paul Pillot, Philippe Le Bris, François-Didier Boyer, Catherine Rameau, Caroline Gutjahr, Alexandre de Saint Germain, Nitzan Shabek

**Affiliations:** Department of Plant Biology, University of California – Davis, Davis, CA 95616; Plant Genetics, TUM School of Life Sciences, Technical University of Munich (TUM), 85354 Freising; Université Paris-Saclay, CEA, CNRS, Institute for Integrative Biology of the Cell (I2BC), 91198, Gif-sur-Yvette, France; Institute of Plant Sciences Paris-Saclay (IPS2), INRAE, CNRS, Université Paris-Sud, Université d’Evry, Université Paris-Diderot, 91405 Orsay, France; Agroécologie, AgroSup Dijon, INRAE, Univ Bourgogne, Univ Bourgogne Franche-Comté, 21000 Dijon, France; Institut Jean-Pierre Bourgin, INRAE, AgroParisTech, Université Paris-Saclay, 78000, Versailles, France; Université Paris-Saclay, CNRS, Institut de Chimie des Substances Naturelles, UPR 2301, 91198, Gif-sur-Yvette, France

## Abstract

KAI2 are plant α/β hydrolase receptors, which perceive smoke-derived butenolide signals (karrikins) and putative endogenous, yet unidentified phytohormones (KAI2-ligands, KLs). The number of functional KAI2 receptors varies among plant species. It has been suggested that *KAI2* gene duplication and sub-functionalization plays an adaptative role for diverse environments or ligand diversification by altering the receptor responsiveness to specific KLs. Legumes represent one of the largest families of flowering plants and contain many essential agronomic crops. Prior to legume diversification, *KAI2* underwent duplication, resulting in *KAI2A* and *KAI2B*. Integrating plant genetics, ligand perception and enzymatic assays, and protein crystallography, we demonstrate that *Pisum sativum* KAI2A and KAI2B act as receptors and enzymes with divergent ligand stereoselectivity. KAI2B has a stronger affinity than KAI2A towards the KAI2-ligand (-)-GR24 and remarkably hydrolyses a broader range of substrates including the strigolactone-like isomer (+)-GR24. We determine the crystal structures of PsKAI2B in apo and butenolide-bound states. The biochemical and structural analyses as well as recorded mass spectra of KAI2s reveal a transient intermediate on the catalytic serine and a stable adduct on the catalytic histidine, further illuminating the role of KAI2 not only as receptors but also as *bona fide* enzymes. Our work uncovers the stereoselectivity of ligand perception and catalysis by evolutionarily diverged KAI2 receptors in KAR/KL signaling pathways and proposes adaptive sensitivity to KAR/KL and strigolactone phytohormones by KAI2B.

## Introduction

Karrikins (KARs) are a family of butenolide small molecules produced from the combustion of vegetation and are bio-active components of smoke (*1–3*). These molecules are capable of inducing germination of numerous plant species, even those not associated with fire or fire-prone environments such as Arabidopsis (*1–6*). Through studies in Arabidopsis, KAR sensitivity was shown to be dependent on three key proteins: a KAR receptor α/β hydrolase KARRIKIN INSENSITIVE2 (KAI2), an F-box MORE AXILLARY GROWTH 2 (MAX2) component of the Skp1-Cullin-F-box (SCF) E3 ubiquitin ligase, and the target of ubiquitination and degradation, the transcriptional corepressor SMAX1/SMXL2 (*7–11*). An increasing number of studies shows that the KAR signaling components are involved in the regulation of a number of plant developmental processes including seedling development, leaf shape, cuticle formation, and root development (*8, 12–15*). Furthermore, they play critical roles in arbuscular mycorrhiza symbiosis and abiotic stress response (*16–18*).

The striking similarities between KAR and strigolactone (SL) signaling pathways have been the focus of an increasing number of studies. Both SLs and KARs share a similar butenolide ring structure but instead of the KAR pyran moiety, the butenolide is connected via an enol ether bridge to either a tricyclic lactone (ABC rings) in canonical SLs, or to a structural variety in non-canonical SLs (*19, 20*). The receptor for SL, DWARF14 (D14) shares a similar α/β hydrolase fold as KAI2 and a parallel signaling cascade requiring the function of the MAX2 ubiquitin ligase and degradation of corepressors (SMXL6, 7 and 8), which belong to the same protein family as SMAX1/SMXL2 (*7, 8, 11, 21*). Unlike KARs, SLs are plant hormones that act endogenously, but were also found to be exuded by plant roots. SLs affect diverse responses such as hyphal branching of arbuscular mycorrhizal (AM) fungi to enhance the efficiency of root colonization, germination of root parasitic plant species, shoot branching, lateral root formation, primary root growth, secondary growth in the stem, leaf senescence, and adventitious root formation (*22–28*). Notably, KAI2 family receptors have undergone numerous duplication events within various land plant lineages. D14 was found to be an ancient duplication in the KAI2 receptor in the seed plant lineage followed by sub-functionalization of the receptor, enabling SL perception (*29–32*). This sub-functionalization has been highlighted by the observations that D14 and KAI2 are not able to complement each other’s functions in planta (*33–37*). While the role of the D14 receptor in SL signaling is better established, KAI2 receptors and KAR signaling are less understood. A central question in receptor diversity has been the evolutionary purpose and functional significance of *KAI2* duplication events, including but not limited to the event that led to the distinct SL receptors D14s. D14s and KAI2s contain over a 70% sequence similarity but confer unique functions in plant. Notably, within the KAI2 family the substitution of a few amino acids within the ligand binding site can alter ligand specificity between KAI2 duplicated copies in *Brassica tournefortii* and *Lotus japonicus* (*35, 38*). Additionally, given that KAR signaling governs diverse developmental processes including those unrelated to fire, KAI2s are thought to perceive endogenous ligands, which remain elusive and tentatively named KAI2-Ligands (KLs) (*29, 30, 37, 39*). Therefore, the ability to alter the specificity of KAI2 receptors to different ligands is likely to be correlated to their ability to perceive distinct KLs. Thus far, several crystal structures of KAI2/D14 receptors have been reported and led to a greater understanding of receptor-ligand perception towards certain ligands (*9, 21, 32, 34, 40–42*). However, the divergence between duplications of KAI2 receptors to confer altered ligand perception and hydrolysis specificities has been only partially addressed for few plant species at the physiological and biochemical level, and a detailed structural examination is still missing (*34–36, 38*).

Legumes represent one of the largest families of flowering plants and contain many essential crops. Beyond their agronomic value, most legume species are unique among plants because of their ability to fix nitrogen by utilizing symbiosis with rhizobia, in addition to AM fungi symbiosis. Because of the potential functional diversification and specialization of KAI2-ligands, in this study, we examined the physiological and biochemical functions of divergent KAI2 receptors in a legume, using *Pisum sativum (Ps)* as a model. We found that *Pisum sativum* expresses three distinct *KAI2* homologues, two of which, *KAI2A* and *KAI2B* have sub-functionalized, while *KAI2C* is a pseudogene. Using comprehensive biochemical characterization, we show that these divergent receptors display distinct ligand sensitivities and hydrolytic activities. We further substantiate these findings *in planta* by investigating the sensitivities to ligands on hypocotyl elongation in Arabidopsis transgenic complementation lines expressing PsKAI2A and PsKAI2B, as well as studying phenotypic effects in *Pisum sativum* wild-type and *kai2* mutants. Strikingly, KAI2B, was more reactive than KAI2A towards SL/KARs stereoisomers. The diverged receptor was able to cleave strigolactone synthetic analog (+)-GR24, although not to the same extend as D14/RMS3, suggesting that PsKAI2B evolved the ability to sense SL-like ligands. To further address this notion, we determine the structure of an evolutionarily diverged PsKAI2B in *apo* and a unique butenolide-bound state at high resolution (1.6 Å and 2.0 Å, respectively). Unlike the D14 *α/β* hydrolase, mass spectrometry analysis and structural examination reveal a mode of ligand perception and hydrolysis by PsKAI2, that involves an intermediate step in which the catalytic serine is transiently bound to a moiety of the ligand and then form a stable adduct with the catalytic histidine. Altogether, in this study we identify and characterize divergent KAI2 receptors, reveal their distinct function and ligand sensitivities and illuminate KAI2s enzymatic mechanism. Better understanding of the evolution of plant *α/β* hydrolase receptors and their functional adaptation in KAR/KL/SL sensing, in particular in a key crop, will have far-reaching impact on the implications of KAR/KL signaling pathways in agro-systems and food security.

## Results

### Identification and characterization of the Pisum sativum KAI2 genes

To characterize the karrikin sensing machinery in legumes, we built a phylogenetic tree of representative legume *KAI2*s. *KAI2* has undergone two independent duplication events in the legume lineage prior to its diversification followed by the loss of KAI2C in the hologalegina clade resulting in distinct KAI2A and KAI2B protein receptors (**Fig. 1a** and **Fig. S1**) (*38*). We focused on the hologalegina representative *Pisum sativum* for which a high-quality, annotated genome sequence has been recently obtained (*43*). We identified three *KAI2* homologs in the pea genome (*43*) that clearly group within the core KAI2 clade by phylogenetic analysis. One (Psat4g083040) renamed *PsKAI2B*, grouped in the same subclade as the legume *KAI2B*s (including *Lotus japonicus, Lj*, *KAI2B* (*38*)), and two Psat2g169960 and Psat3g014200 respectively termed *PsKAI2A* and *PsKAI2C*, in the same subclade as the legume *KAI2A*s (including *LjKAI2A* (*38*)) (**Fig. 1a** and **Fig. S1**). *PsKAI2C* was detected by PCR in the genomic, but not the cDNA, and appear to be pseudogene because the predicted encoded protein is truncated at 128 amino acids (aa) due to a premature stop codon at 387 nucleotides after the translation initiation site (**Fig. 1b**). By cloning the *PsKAI2A* coding sequence (CDS) we identified two transcripts for this gene, corresponding to two splice variants (**Fig. 1b** **and Fig. S2**). The transcript PsKAI2A.1 results from intron splicing and encodes a protein of 305 aa. Thus, this protein shows a C-terminal extension of 33 aa similar to LjKAI2A (**Fig. S2**), missing in other KAI2 proteins. The PsKAI2A.2 transcript arises from the intron retention and shows a premature STOP codon two nucleotides after the end of the first exon. This leads to a 272 aa protein representing a similar size to other KAI2s described (**Fig. 1b** and **Fig. S2**). Due to the similar size, lack of introns, and evolutionary conservation of the residues contained in the PsKAI2A.2 protein, PsKAI2A hereafter will refer to the PsKAI2A.2 protein. From this analysis, it is clear that the KAI2 clade has undergone an independent duplication event in the legume lineage, resulting in two functional KAI2A and KAI2B forms (**Fig. S1a-b**). To examine potential functional divergence between the PsKAI2A.2 and PsKAI2B forms, we first analyzed the aa sequences and identified notable alterations in key residues, of which numerous are likely to be functional changes as indicated in later analyses (**Fig. S3**). To further characterize divergence of these genes, we studied their expression patterns in various tissues of the *Pisum* plant (**Fig. 1c-d**). Interestingly, *PsKAI2A* was ten-fold more highly expressed in comparison to *PsKAI2B* and the expression in the roots differed between the two forms, suggesting sub-functionalization between transcriptional regulatory sequences of *PsKAI2A* and *PsKAI2B*.

**Fig. 1.**
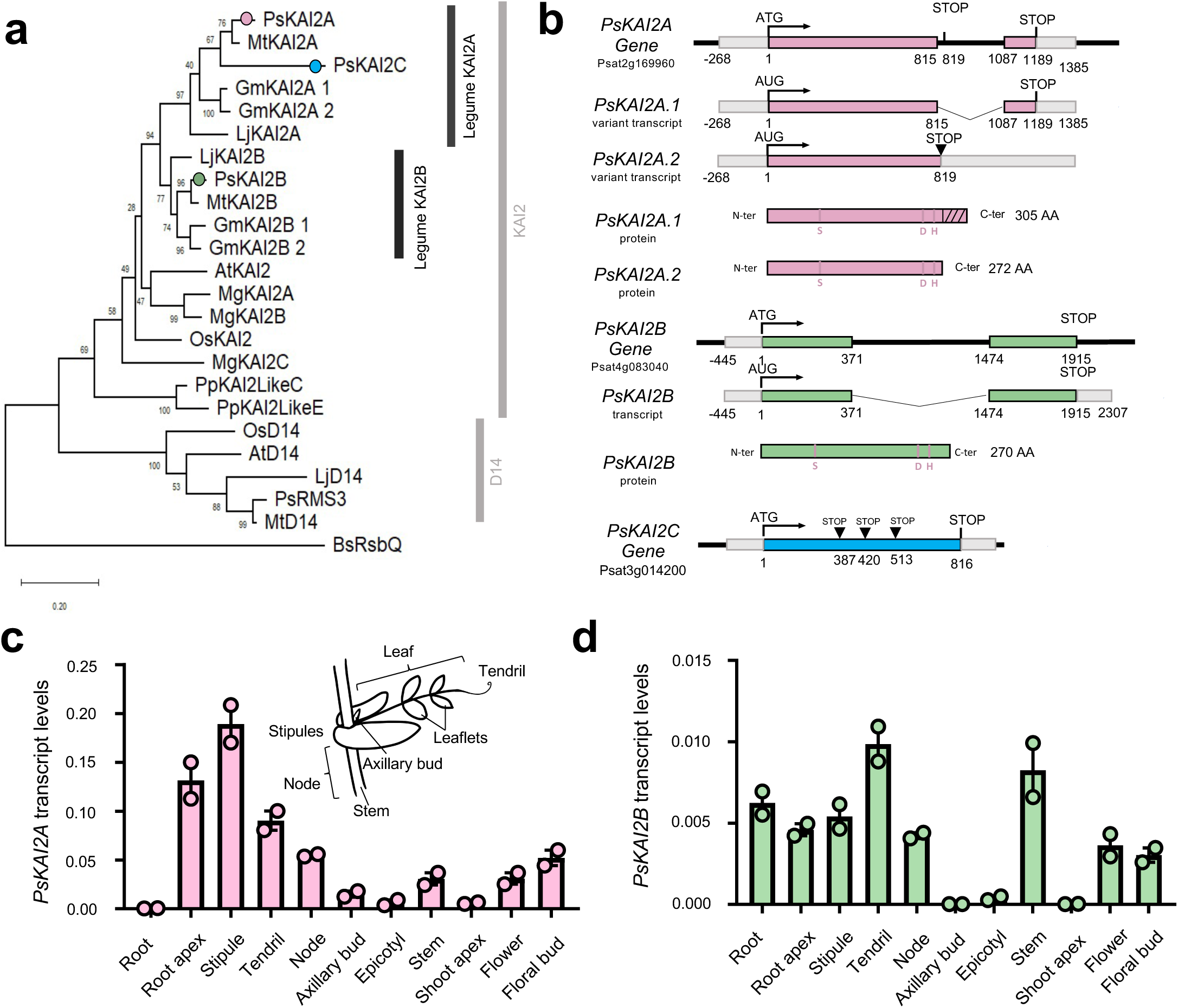
Evolutionary analysis and differential expression of the legume Pisum sativum KAI2s. **(a)** Maximum likelihood phylogeny of 24 representative KAI2 amino acid sequences. Node values represent percentage of trees in which the associated taxa clustered together. Vertical rectangles highlight distinct KAI2 family clades. Black circle indicates legume duplication event. Pink, green, and blue circles mark the position of PsKAI2As, PsKAI2B, and PsKAI2C respectively. The tree is drawn to scale, with branch lengths measured in the number of substitutions per site. **(b)** PsKAI2A, PsKAI2B and PsKAI2C are homologues to AtKAI2 and encode α-β/hydrolases. Schematic representation of the PsKAI2A, PsKAI2B and PsKAI2C genes; Exons are in pink, green and blue lines, non-coding sequences colored in thin black lines and UTR regions shown as thick gray lines. Bases are numbered from the start codon. PsKAI2A shows 2 splicing variants. Spliced introns are shown as bent (“V”) lines. Inverted triangle (▾) indicates premature termination codons. Catalytic triad residues are indicated in pink. The hatched part indicated the C-terminus extension of the PsKAI2.1 protein (**c-d**) Differential expression pattern of PsKAI2A (**c**, pink) and PsKAI2B (**d**, green). Transcript levels in the different tissues of 21 old wild-type Pisum sativum plants (cv. Terese) were determined by real-time PCR, relative to PsEF1α. Data are means ± SE (n = 2 pools of 8 plants). Inset drawing of a node showing the different parts of the pea compound leaf.

### Identification and phenotypic examination of Pskai2a and Pskai2b TILLING mutants

To investigate the function of *PsKAIA* and *PsKAI2B*, mutants in both genes were identified via Targeting-Induced Local Lesions IN Genomes (TILLING) using the mutagenized Caméor population (*44, 45*). Twenty mutations in *PsKAI2A* and sixteen mutations is *PsKAI2B* were identified (**Table S1**). Among them, five of the *PsKAI2A* and three of the *PsKAI2B* mutations were predicted as non-synonymous and may result in mutated amino acids that compromise the protein function (**Table S1,** **Fig. 2a, Fig. S4**). The comparison between wild-type and *Pskai2* single mutants revealed no detectable shoot architecture differences in either branch number or in plant height. The double mutant *Pskai2a-6 Pskai2b-3* showed a reduce height and decreased branching (**Fig. S5a-c).** Similar to the other legume *L. japonicus* and in contrast to *Arabidopsis thaliana* (*38, 46*), the root hair length of the double mutant *Pskai2a-6 Pskai2b-3* in pea is not significantly different from wild-type. Together, under our growth conditions, KAI2 requirement for root hair elongation differs among plant species/families and may be absent from legumes. (**Fig. S5d, e**).

**Fig. 2.**
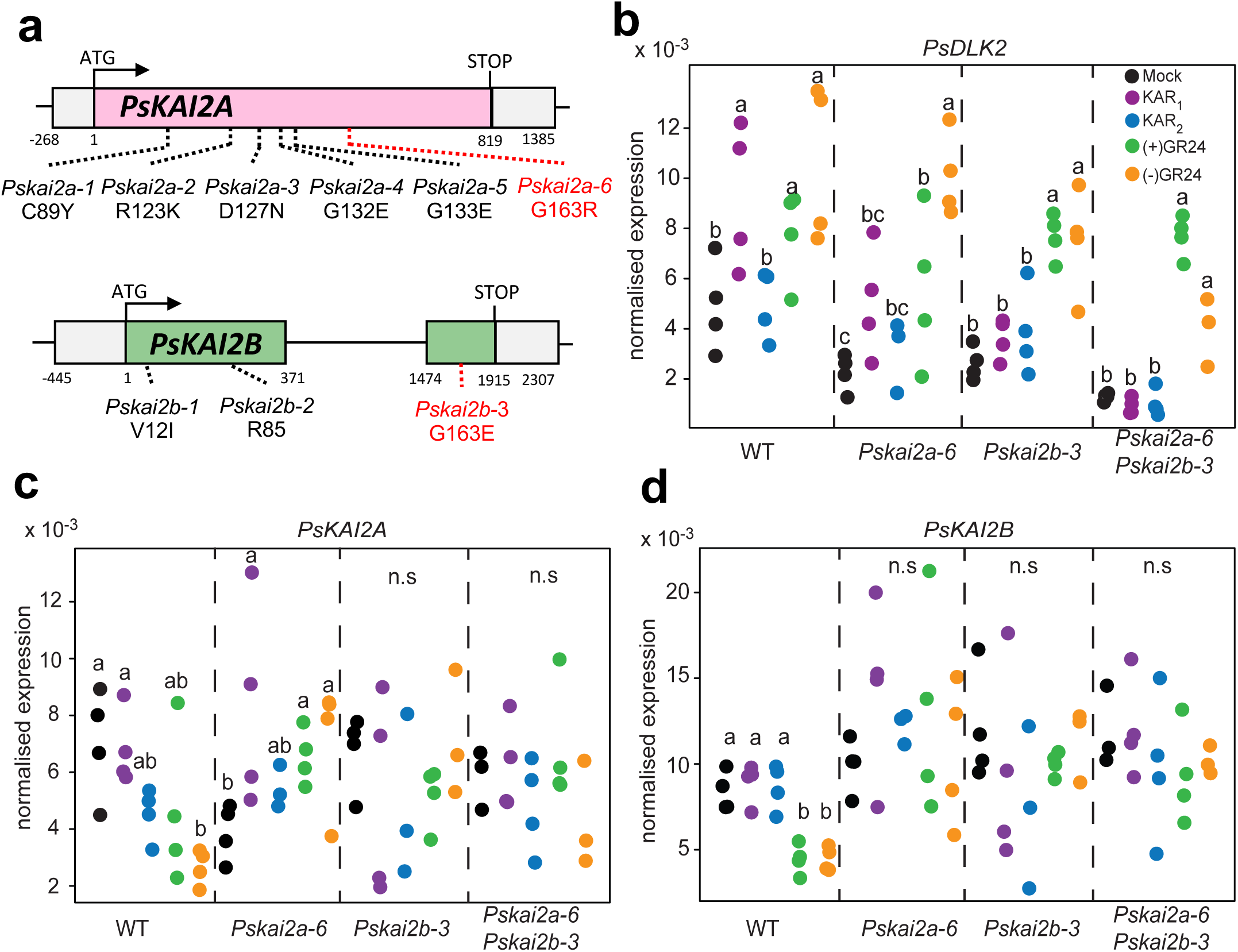
Characterization of the Pskai2 mutants. **(a)** Gene structure of PsKAI2 and locations of mutations. Bases are numbered from the start codon. Point mutations are indicated by dotted lines (black and red for the one studied here) (**b-d**) RT-qPCR-based expression of PsDLK2 (**b**), PsKAI2A (**c**), PsKAI2B (**d**) in roots of 10-day old P. sativum plants after 4 hours treatment with solvent (Mock) or 3 μM KAR1 or (+)-GR24 or (–)-GR24. Expression values were normalized to those of the housekeeping gene TUBULIN (n = 3–4). Letters indicate significant differences versus mock treatment (Kruskal-Wallis Test, p<0.05).

### Karrikin signaling reporter DLK2 expression is mediated by KAI2s in P. sativum roots in a ligand-specific manner

It was recently shown, in *L. japonicus* that the root system architecture can be modulated by KAR_1_ but not by KAR_2_ treatment, and the expression of the KAR signaling marker gene, *DLK2* in roots, was responsive only to KAR_1_ but not to KAR_2_ (*38*). Furthermore, LjKAI2A and LjKAI2B have distinct ligand-binding specificities since LjKAI2A but not LjKAI2B can perceive (–)-GR24 *in vitro*. This alteration in perception depends on the divergent amino acid F157/W158 within the ligand binding pocket, and indeed in roots, LjKAI2A but not LjKAI2B can induce *DLK2* expression in response to (–)-GR24 (*38*). Intriguingly, because F157/W158 amino acid difference between KAI2A and KAI2B is not conserved across most legumes, this raises the question how pea roots respond to the range of artificial ligands. Thus, we treated pea wild-type, *kai2a-6, kai2b-3, and kai2a-6 kai2b-3* double mutant roots with KAR_1_, KAR_2_, (+)-GR24, and (–)-GR24 and performed RT-qPCR analysis of *PsDLK2* transcript accumulation (**Fig. 2b**). Similar to *L. japonicus*, *DLK2* was induced in pea wild-type roots only in response to KAR_1_ but not to KAR_2_. Interestingly, without treatment, *PsDLK2* expression was significantly reduced in both single mutants (*kai2a-6* and *kai2b-3*) indicating that *KAI2A* and *KAI2B* are not fully redundant and even more so in the double mutant (*kai2a-6 kai2b-3*) (**Fig. 2b**). This expression pattern strongly suggests that PsKAI2A and PsKAI2B functionally mediate KAR signaling and validates that the *kai2a-6* and *kai2b-3* mutants perturb protein function. Surprisingly, in response to KAR_1,_ *Pskai2a-6* shows only a minor induction of *PsDLK2,* whereas *Pskai2b-3* shows no *PsDLK2* induction. Given that both *PsKAI2A* and *PsKAI2B* transcripts accumulated to similar levels (**Fig. 2c, d**), these data suggest that either PsKAI2B protein accumulate to higher levels than PsKAI2A or it is more active in KAR_1_ perception. In all genotypes, *PsDLK2* was induced by (–)-GR24 treatment, which is consistent with previous studies in Arabidopsis where (–)-GR24 also acts as ligand for the SL receptor D14 (*46, 47*). Unlike the observations with *L. japonicus* (*38*), all pea genotypes showed a significant induction of *DLK2* expression in response to (–)-GR24, with a significantly stronger induction in the wild-type and the single mutants as compared to the double mutant. Thus, in pea, (–)-GR24 is not only perceived by KAI2 but to some extent also by the SL receptor D14, similar to previous observations in Arabidopsis (*46*). However, we cannot exclude the possibility that the mutated proteins are still able to perceive (–)-GR24, even though they lost sensitivity to KAR_1_.

### PsKAI2A, but not PsKAI2B, rescues inhibition of hypocotyl elongation in Arabidopsis kai2 mutants

Since we could not distinguish a possible differential sensitivity of PsKAI2A and PsKAI2B in the pea background (as previously observed in *L. japonicus* (*38*)), we performed a cross-species complementation by transforming the Arabidopsis *htl-3* (also known as *kai2*) mutant with *PsKAI2A.2* and *PsKAI2B* (**Fig. 3**). The genes were expressed as fusions with *mCitrine* or *GUS* and driven by the native *AtKAI2* promoter (p*AtKAI2*) (**Fig. S6b-c**). To test protein functionality we performed the widely used hypocotyl elongation assay (*33, 39*) under low light conditions, which causes an elongated hypocotyl phenotype of the *htl-3* mutant when compared to the wild-type Columbia (Col-0). Remarkably, all transgenes, except *PsKAI2B*, completely or partially restored hypocotyl length of *htl-3* to the wild-type length (**Fig. 3**). Also interestingly, all lines (even *htl-3*), except those complemented with PsKAI2A responded to (-)-GR24 (**Fig. 3**). We repeated the complementation for the *kai2-2* mutant in the Landsberg erecta (Ler) background. The data confirmed that the two *PsKAI2A* splice forms restored the reduced hypocotyl length in *kai2-2* but the proteins did not mediate responses to (–)-GR24, whereas PsKAI2B did not restore the wild-type hypocotyl length but mediated a response to (–)-GR24 (**Fig. S6**). Interestingly, these results suggest that PsKAI2A can perceive endogenous KL in Arabidopsis but does not perceive the synthetic (-)-GR24. In contrast, PsKAI2B is unable to perceive KL but is sensitive to (-)-GR24. These results propose that *PsKAI2A* is the functional orthologue of Arabidopsis *KAI2* while PsKAI2B has diverged and adapted to perceive different ligands.

**Fig. 3.**
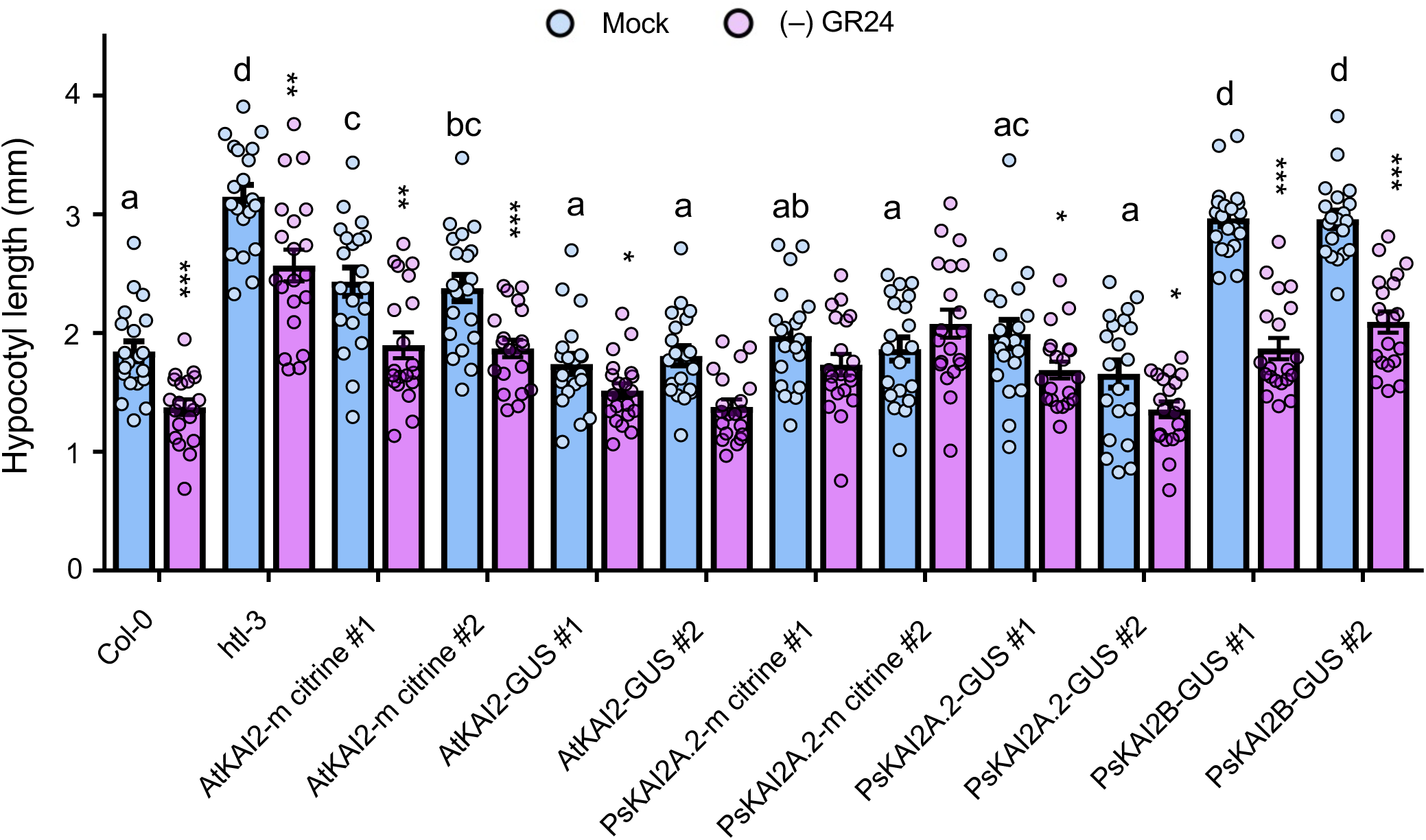
Arabidopsis hypocotyl elongation complementation assays with PsKAI2s. Hypocotyl length of 7-day-old seedlings grown under low light at 21 °C. Data are means ± SE (n = 20-24; 2 plates of 10-12 seedlings per plate). Light blue bars: Mock (DMSO), lavender bars: (–)-GR24 (1µM). Complementation assays using the AtKAI2 promoter to express AtKAI2 (control) or PsKAI2 genes in the htl-3 mutant background (Ler ecotype) as noted above the graph. Proteins were tagged with mCitrine or GUS protein. For DMSO controls, statistical differences were determined using a one-way ANOVA with a Tukey multiple comparison of means post-hoc test, statistical differences of P<0.05 are represented by different letters. Means with asterisks indicate significant inhibition compared to mock-treated seedlings with *** corresponding to p ≤ 0.001 and * to p ≤ 0.01, as measured by t-test.

### Altered ligand binding specificity and activity between PsKAI2s

To further investigate the distinct ligand selectivity, we purified PsKAI2 recombinant proteins and studied their ligand-interaction and ligand-enzymatic activities using various complementary assays (**Fig. 4-5** and **Fig. S7-S9**). We first examined PsKAI2A.2 and PsKAI2B ligand interactions via the thermal shift assay (Differential Scanning Fluorimetry, DSF) with various KAI2/D14 family ligands including (+) and (–)-GR24 enantiomers (also known as GR24^5DS^ and GR24*^ent^*^-5DS^, respectively), and (+)- and (–)-2’-*epi*-GR24 (also known as GR24*^ent^*^-5DO^ and GR24^5DO^, respectively) (*47*) (**Fig. 4c-f, Fig. S8**). DSF analyses revealed an increased shift in stability in the presence of (–)-GR24 for PsKAI2B as compared to PsKAI2A which has little to no alteration (**Fig. 4c-f**). The other ligands and enantiomers induce no detectable shift in stability for either receptor. An extensive interaction screen using intrinsic fluorescence confirmed that only the (–)-GR24 stereoisomer interacts with PsKAI2 proteins (**Fig. 4b** and **Fig. S9**), and further corroborating the results of the Arabidopsis hypocotyl elongation and the DSF assays. The calculated *K*_d_ shows a higher affinity for (–)-GR24 for PsKAI2B (*K*_d_ = 89.43 ± 12.13 μM) than PsKAI2A (*K*_d_ = 115.40 ± 9.87 μM).

**Fig. 4.**
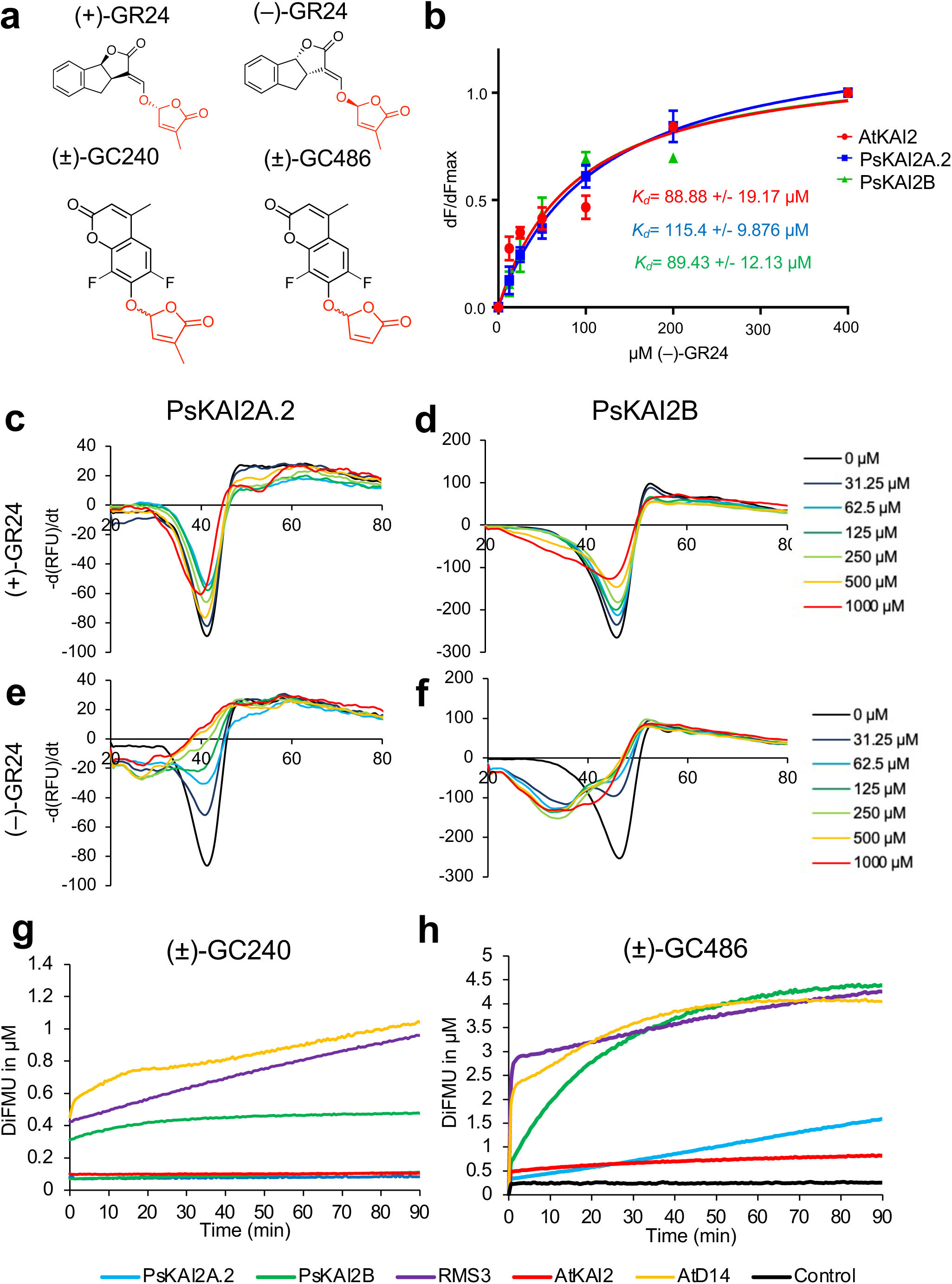
Biochemical analysis of PsKAI2A.2 and PsKAI2B interactions with different GR24 isomers and enzymatic property. **(a)** Chemical structure of ligands used. (**b**) Plots of fluorescence intensity versus SL concentrations. The change in intrinsic fluorescence of AtKAI2, PsKAI2A.2 and PsKAI2B was monitored (see **Fig. S4**) and used to determine the apparent Kd values. The plots represent the mean of two replicates and the experiments were repeated at least three times. The analysis was performed with GraphPad Prism 7.05 Software. (**c-f**) DSF assay. The melting temperature curves of 10 µM PsKAI2A.2 (**c, e**) or PsKAI2B (**d, f**) with (+)-GR24 (c-d), (–)-GR24 (**e-f**) at varying concentrations are shown as assessed by DSF. Each line represents the average protein melt curve for three technical replicates; the experiment was carried out twice. (**g-h**) Enzymatic kinetics for AtD14, AtKAI2, RMS3, PsKAI2A.2 and PsKAI2B proteins incubated with (±)-GC240 (g) or (±)-GC486 (**h**). Progress curves during probes hydrolysis, monitored (λem 460 nm) at 25 °C. Protein catalyzed hydrolysis with 400 nM µM of protein and 20 µM of probes. These traces represent one of the three replicates, and the experiments were repeated at least twice.

**Fig. 5.**
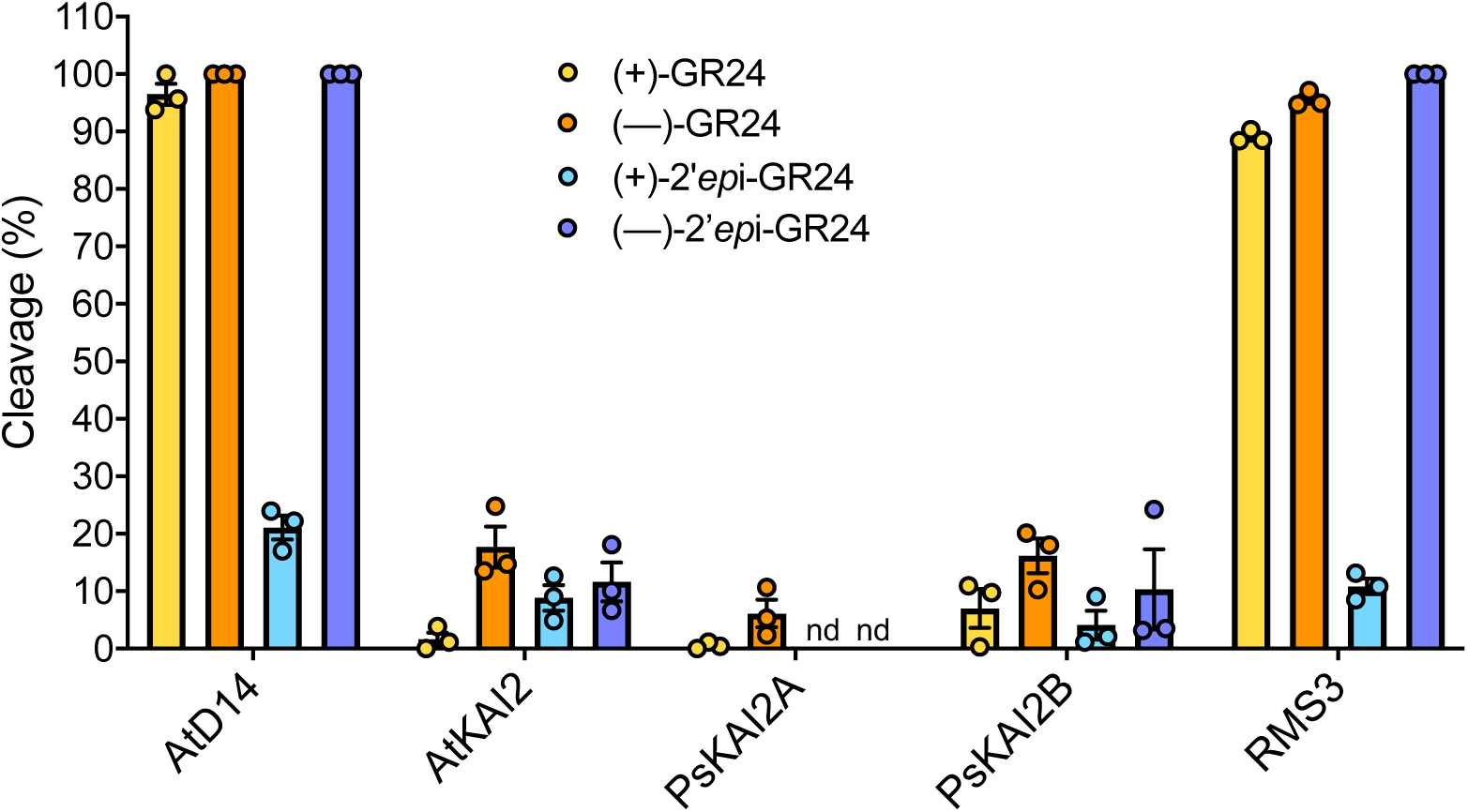
Comparative enzymatic activity of AtD14, AtKAI2, RMS3, PsKAI2A and PsKAI2B proteins with GR24 isomers. UPLC-UV (260 nm) analysis showing the formation of the ABC tricycle from GR24 isomers. The enzymes (10 μM) hydrolysis activity was monitored after incubation with 10 μM (+)-GR24 (yellow), (–)-GR24 (orange), (+)-2’-epi-GR24 (blue), or (–)-2’-epi-GR24 (purple). The indicated percentage corresponds to the hydrolysis rate calculated from the remaining GR24 isomer, quantified in comparison with indanol as an internal standard. Data are means ± SE (n = 3). nd = no cleavage detected.

Because the structural similarity between KAI2s and D14s α/β hydrolases in particular the conserved serine catalytic triad signature of these receptors, we examine the potential catalytic function of KAI2s. The hydrolytic activity of the PsKAI2 proteins towards distinct ligands was quantified in comparison to AtD14, AtKAI2, and RMS3. The proteins were incubated with (+)- GR24, (–)-GR24, (+)-2’-*epi*-GR24 and (–)-2’-*epi*-GR24 in presence of 1-indanol as an internal standard, followed by ultraperformance liquid chromatography (UHPLC)/UV DAD analysis (**Fig. 5**). Interestingly, PsKAI2A can only cleave (–)-GR24, while PsKAI2B is able to cleave (+)-GR24, (–)-GR24 and (–)-2’-*epi*-GR24 stereoisomers. Among the tested ligands, all enzymes show reduced activity towards (+)-2’-*epi*-GR24, and among the KAI2s, PsKAI2A had no detectable activity for (+)-2’-*epi*-GR24. These results strongly indicate that PsKAI2s have distinct stereoselectivity, and PsKAI2B appears to be more reactive. To further investigate the hydrolysis kinetics of PsKAI2 proteins, we performed an enzymatic assay with pro-fluorescent probes that were previously designed for detecting SL hydrolysis (*36, 48*) (**Fig. 4g-h**). As expected, PsKAI2A shows no activity towards the probe (±)-GC240, similar to AtKAI2, as previously reported (*48*). Strikingly, PsKAI2B is able to cleave (±)-GC240, suggesting yet again that unlike PsKAI2A, PsKAI2B has a ligand stereoselectivity that is similar to AtD14 and RMS3 (**Fig. 4g**). Since probes without a methyl group, such as dYLG, can serve as the hydrolysis substrates for AtKAI2 (*49*), we tested the activity of KAI2s and D14s using the (±)-GC486 probe bearing no methyl on D-ring. Notably, PsKAI2B, RMS3, and AtD14 were able to effectively hydrolyze (±)-GC486, whereas PsKAI2A and AtKAI2 showed little activity (**Fig. 4h**). Nonetheless, PsKAI2A and AtKAI2 also exhibit a biphasic time course of fluorescence, consisting of an initial phase, followed by a plateau phase. Comparative analysis of the kinetic profiles shows that PsKAI2B, RMS3 and AtD14 display a higher plateau (1 µM versus 0.3 µM of DiFMU, **Fig. 4h**). Taken together the comparative kinetic analysis substantiates the distinct function of PsKAI2B compared to PsKAI2A not only highlights the unique similarity of PsKAI2B to SL receptors (*8, 10, 36, 40–42, 48, 50*) rather than karrikin receptors (*8–10, 13, 34–36, 38, 49*), but may also offer new insights into the missing link between Karrikin and SL by diverged KAI2s.

### Structural insights into divergence of legume KAI2A and KAI2B

To elucidate the structural basis for the differential ligand selectivity between KAI2A and KAI2B, we determined the crystal structure of PsKAI2B at 1.6 Å resolution (**Fig. 6** and **Table S2**). The PsKAI2B structure shares the canonical *α*/*β* hydrolase fold and comprises base and lid domains (**Fig. 6a**). The core domain contains seven-stranded mixed β-sheets (β1–β7), five α-helices (αA, αB, αC, αE and αF) and five 3_10_ helices (ŋ1, ŋ2, ŋ3, ŋ4, and ŋ5). The helical lid domain (residues 124–195, **Fig. S3**) is positioned between strands β6 and β7 and forms two parallel layers of V-shaped helices (αD1-4) that create a deep pocket area adjoining the conserved catalytic Ser-His-Asp triad site (**Fig. 6b** and **Fig. S3**). Despite the sequence variation (77% similarity between PsKAI2B and AtKAI2, **Fig. S3**), we did not observe major structural rearrangements between PsKAI2B and the previously determined Arabidopsis KAI2 structure (*51*) as shown by an Root Mean Squared Deviation (RMSD) of 0.35 Å for superposition of backbone atoms (**Fig. 6b**). Nonetheless, further structural comparative analyses have identified two residue alterations in positions 129 and 147 within the lid domain which appear to slightly alter the backbone atoms and generally distinguish legume KAI2 proteins from KAI2s in other species (**Fig. S3** and **Fig. 6b**). The asparagine residue in position 129 is more unique to legume KAI2As, and the legume alanine or serine in position 147 has diverged from bulky polar residues compared to other KAI2s. Therefore, it is likely that these amino acids variations play role in downstream events rather than directly modulate distinct ligand perception.

**Fig. 6.**
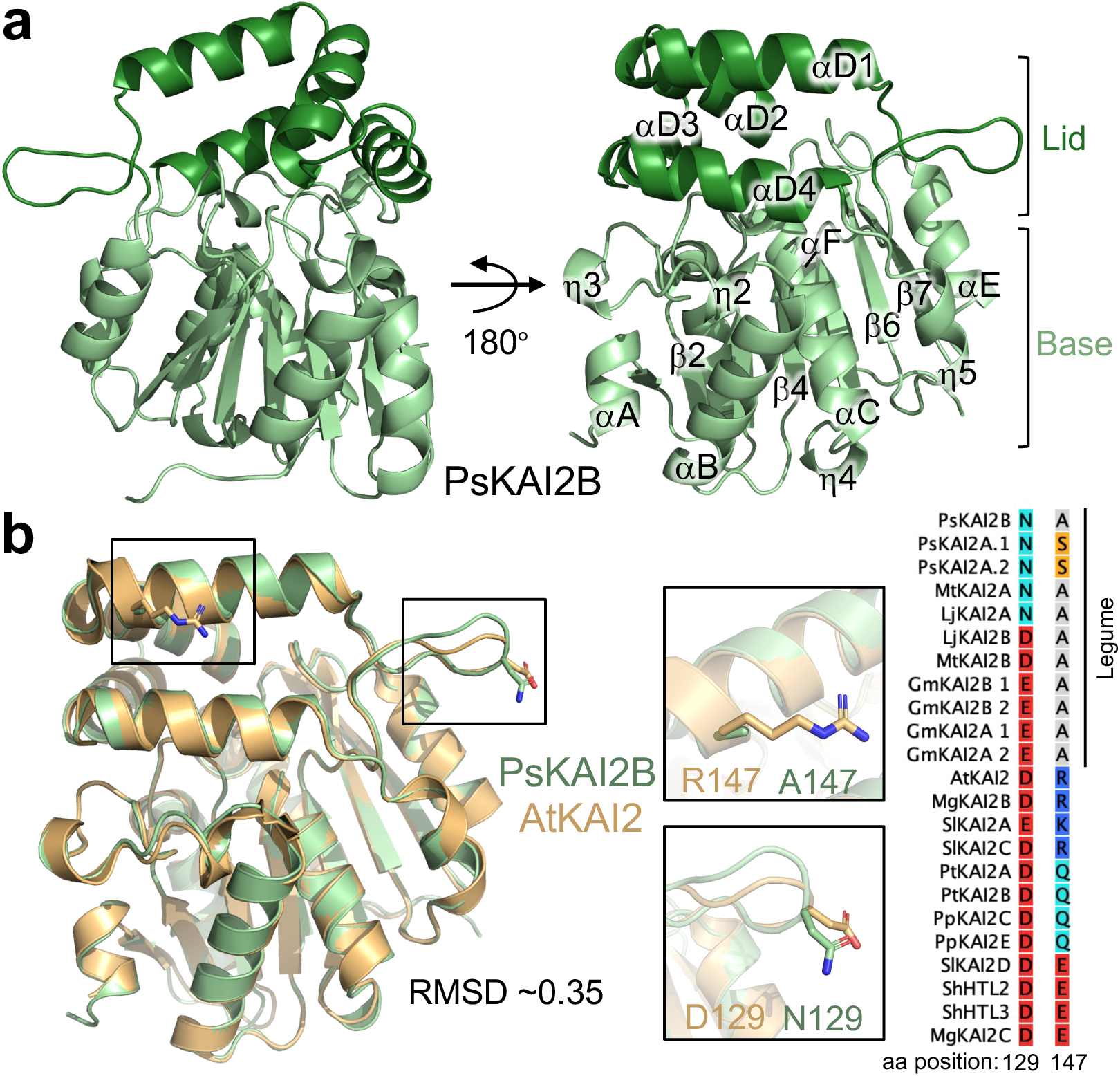
The crystal structure of legume KAI2. (a) Overview of PsKAI2B structure. Lid and base domains are colored in forest and light green respectively with secondary structure elements labeled. (**b**) Structural alignment of PsKAI2B and AtKAI2 (PDB ID: 4HTA) shown in light green and wheat colors respectively. Root-mean-square deviation (RMSD) value of the aligned structures is shown. The location and conservation of legume KAI2 unique residues, alanine in position 147 (A147) and asparagine N129, are highlighted on the structure shown as sticks as well as in reduced Multiple Sequence Alignment from **Fig. S1**.

To further determine the differential ligand specificity between PsKAI2A and PsKAI2B, we utilized the PsKAI2B crystal structure reported here to generate a high probability 3D model for PsKAI2A. As expected, PsKAI2A structure exhibits a similar backbone atom arrangement (RMSD of 0.34 Å) that parallels the PsKAI2B structure (**Fig. 7a**). Nonetheless, we identified eight significant divergent amino acids between the two structures including residues involved in forming the ligand binding pocket as well as solvent-exposed regions (**Fig. 7b-d** and **Fig. S10a-b**). Because these variants are evolutionarily conserved across legumes, the analysis of the underlined residues not only distinguishes between KAI2A and KAI2B in *Pisum* but can be extrapolated to other legume KAI2A/B diverged proteins. Structural comparative analysis within the ligand-binding pocket shows divergent solvent accessibility between PsKAI2A and PsKAI2B (**Fig. 7b**). PsKAI2B exhibits a structural arrangement that results in a larger volume of the hydrophobic pocket (125.4 A^3^) yet with a smaller entrance circumference (30.3 Å) than PsKAI2A (114.8 A^3^ and 33.6 Å, respectively, **Fig. 7b**). Further *in silico* docking experiments of (–)-GR24 with PsKAI2B results in a successful docking of the ligand that is totally buried in the pocket and positioned in a pre-hydrolysis orientation nearby the catalytic triad. In contrast, docking experiments of (–)-GR24 with PsKAI2A results in more restricted interaction where the ligand is partially outside the pocket (**Fig. S10c)**. Notably, there are five key residues that are found to directly alter the pocket morphology (**Fig. 7c** and **Fig. S10a-b**). Among these residues, L160/S190/M218 in PsKAI2A and the corresponding residues, M160/L190/L218 in PsKAI2B are of particular interest because of their functional implications in the pocket volume and solvent accessibility (**Fig. 7d**). Interestingly, the variant in position 218 places it in the center of the Asp loop (D-loop, region between β 7 and αE, **Fig. 7c-d**), that has been recently suggested to impact D14 protein-protein interactions in SL signaling (*52, 53*). Residues 160 and 190 are of great interest because of their direct effect on ligand accessibility and binding pocket size. Residue 160 is positioned at the entrance of the ligand-binding pocket in helix αD2, thus the substitution of leucine (L160 in KAI2A) to methionine (M160 in KAI2B) results in modifying the circumference of PsKAI2B pocket entrance (**Fig. 7b-d**). While both L160 and M160 represent aliphatic non-polar residues, the relative low hydrophobicity of methionine as well as its higher plasticity are likely to play a major role in modifying the ligand pocket. The conserved divergence in residue 190 (S190 in PsKAI2A and L190 in PsKAI2B, **Fig. 7d**) is positioned in helix αD4 and represents a major structural arrangement at the back of the ligand envelope. Because leucine has moderate flexibility compared to serine and much higher local hydrophobicity, this variation largely attributes to the changes in the pocket volume as well as fine-tunes available ligand orientations.

**Fig. 7.**
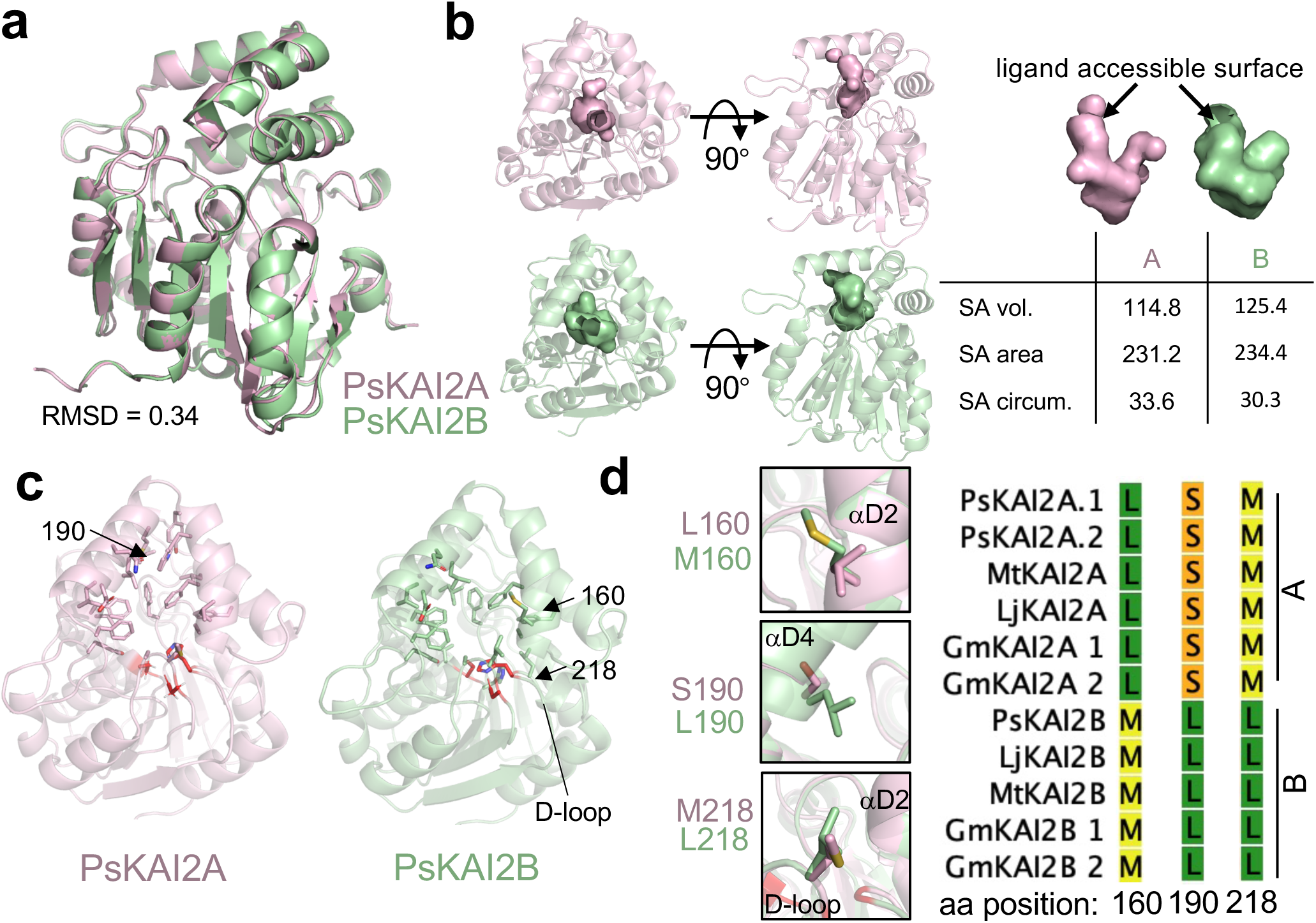
Structural divergence analysis of legume KAI2A and KAI2B. **(a)** Structural alignment of PsKAI2A and PsKAI2B shown in pink and light green colors respectively. RMSD of aligned structures is shown. (**b**) Analysis of PsKAI2A and PsKAI2B pocket volume, area, and morphology is shown by solvent accessible surface presentation. Pocket size values were calculated via the CASTp server. (**c**) Residues involved in defining ligand-binding pocket are shown on each structure as sticks. Catalytic triad is shown in red. (**d**) Residues L/M160, S/L190, and M/L218 are highlighted as divergent legume KAI2 residues, conserved among all legume KAI2A or KAI2B sequences as shown in reduced Multiple Sequence Alignment from **Fig. S1**.

To further examine whether the diverged residues directly impact ligand perception. We produced the recombinant reciprocal swap mutants PsKAI2A.2^L160M, S190L^ and PsKAI2B^M160L, L190S^, and tested their sensitivity to selected ligands by DSF. Remarkably, the swap of PsKAI2A.2 and PsKAI2B at residues 160 and 190 resulted in a loss of sensitivity of PsKAI2B^M160L, L190S^ towards (–)-GR24 ligand, yet with no gain of sensitivity of PsKAI2A.2^L160M, S190L^ (**Fig. S11**). This demonstrates that the variation in residues 160 and 190 in KAI2s are necessary for ligand perception and selectivity but may not be entirely sufficient for PsKAI2B perception of (–)-GR24. Notably, structural analysis of the ligand-envelopes of PsKAI2A.2^L160M, S190L^ and PsKAI2B^M160L, L190S^ in comparison to their WT counterparts further corroborates the adapted pocket size in PsKAI2B^M160L, L190S^, which explain the observed loss of sensitivity to (–)-GR24 (**Fig. S11e**).

### Structural and functional elucidation of ligand hydrolysis mechanism by PsKAI2B receptor

To examine the molecular interaction of PsKAI2B with the enantiomer (–)-GR24, we co-crystallized and determined the structure of PsKAI2B-(–)-GR24 at 2.0 Å resolution (**Fig. 8a** and **Table S2**). Electron density map analysis of the ligand-binding pocket revealed the existence of a unique ring-shaped occupancy that is contiguously linked to the catalytic serine (S95) (**Fig. 8a-b**). The structural comparison of the backbone atoms between apo-PsKAI2B and PsKAI2B- (–)-GR24 did not reveal significant differences (**Fig. S12a**) and is in agreement with previously reported *apo* and ligand bound D14/KAI2 crystal structures (*9, 10, 21, 32, 41*). This similarity suggests that a major conformational change may indeed occur as proposed for D14 (*53*). It may happen after the nucleophilic attack of the catalytic serine and the (–)-GR24 cleavage which is likely to be a highly unstable state for crystal lattice formation. Further analysis suggests that the 5-hydroxy-3-methylbutenolide (D-OH ring), resulting from the (–)-GR24 cleavage, is trapped in the catalytic site (**Fig. S12b-d)**. The lack of a defined electron density fitting with the tricyclic lactone (ABC ring) may exclude the presence of the intact GR24 molecule. Other compounds present in the crystallization condition were tested for their ability to occupy the S95-contiguous density, and the D-OH group of (–)-GR24 demonstrated the highest calculated correlation coefficient (CC) score and the best fit in the PsKAI2B co-crystal structure (**Fig. S12c**). Additional tests of D-OH binding including *in silico* docking simulations and analyses revealed a high affinity for D-OH in a specific orientation and in agreement with the structure presented here (**Fig. S12d**). The most probable orientation of the D-OH positions the methyl group (C4’) together with the hydroxyl group of D-OH towards the very bottom/back of the pocket near the catalytic serine, where the O5” atom is coordinated by both N atoms of F26 and V96 (**Fig. 8b-c**). The hemiacetal group (C2’) of D-OH is oriented towards the access groove of the pocket with angles (between carbon and oxygen atoms) supporting the captured D-OH in an orientation in which cleavage of the intact (–)-GR24 may have taken place. The C5’ of D-OH appears to form a covalent bond with O*γ* of S95 (dark gray line in **Fig. 8c**) and generates a tetrahedral carbon atom. The overall positioning of this molecule is strictly coordinated by F26, H246, G25, and I193 residues. Notably, the electron density around the S95 does not display an open D-OH group (2,4,4,-trihydroxy-3-methyl-3-butenal as previously described for OsD14 (*21*)) that could directly result from the nucleophilic attack event, but rather more closely corresponds to a cyclized D-OH ring linked to the S95. The D-OH ring is likely to be formed by water addition to the carbonyl group at C2’ that is generated after cleavage of the enol function and cyclization to re-form the butenolide (**Fig. 8d**). Based on the suggested mechanism of SL hydrolysis by D14 (*36, 48, 50, 53*), we propose that the formation of the S95-adduct serves as a highly transient intermediate before its transfer to the catalytic histidine residue. Therefore, to further examine (–)-GR24 catalysis by PsKAI2s, we recorded mass spectrometry (MS) spectra under denaturing conditions with PsKAI2B and PsKAI2A. As expected, a mass shift occurred corresponding to an intermediate covalently bound to PsKAI2s (**Fig S13a-d**). Following digestion, a peptide with additional mass of 96 Da was detected on the catalytic residue H246, further corroborating the transient nature of S95-adduct that was capture in the crystal structure (**Fig. S13e-f**). Collectively, the crystal structure of PsKAI2B bound to GR24 and the MS data show transient and stable intermediates attached to S95 and H246 of the catalytic triad and reveal the mode of action of KAI2 as receptors and enzymes (**Fig. 8d**).

**Fig. 8.**
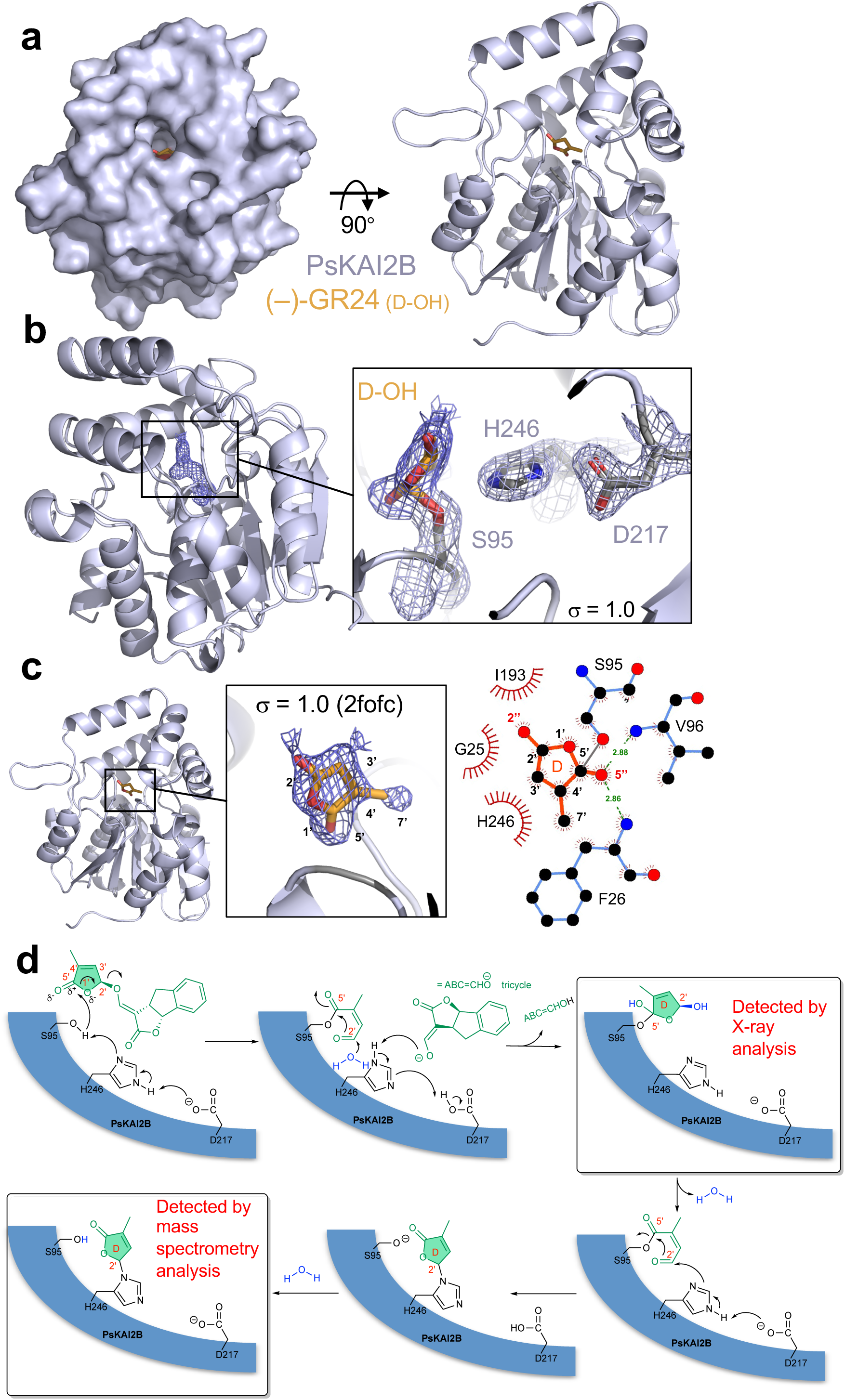
Structural basis of PsKAI2B ligand interaction. **(a)** Surface (left) and cartoon (right) representations of PsKAI2B crystal structure in complex with (–)-GR24 D-OH ring. Protein structure is shown in blue/gray and ligand in orange. (**b**) Close-up view on ligand interactions and contiguous density with the catalytic serine S95. Electron density for the ligand is shown in navy blue and blue/gray mesh for the labeled catalytic triad. The contiguous density between S95 and the D-OH ring indicates a covalent bond. The electron density is derived from 2mFoDFc (2fofc) map contoured at 1.0s. (**c**) Side view of PsKAI2B-D-OH structure shown in cartoon with highlighted (orange) the intact D-OH ring structure. 2-D ligand interaction plot was generated using LigPlot+ software. Dark grey line represents S95-D-OH ring covalent bond. (**d**) Schematic diagram of the proposed mechanism for the formation of the D-ring intermediate covalently bound to S95 in a first time and then to H246.

## Discussion

The coevolution between receptors and ligands in diverse environments throughout plant evolution is of wide interest in many biological fields. In particular, characterization of the emerging karrikin/KL signaling in non-fire following plants has been of increasing importance in plant signaling at large. While there are many missing pieces in the karrikin/KL signaling puzzle, it is clear that KAI2 is the key sensor in this pathway(s). The striking evolutionary conservation of KAI2 receptors in all land plants is not fully explained by the limited natural occurrence of smoke-derived karrikin molecules as well as non-fire following species. Also, an increasing number of studies suggests that the functions of KAI2s are preserved to regulate plant development and response to stresses by perceiving KL signals from either internal or external sources. Most importantly, the intriguing potential dual function of KAI2s as receptors and enzymes remained largely elusive. Here, we identified and characterized the KAI2 receptors in pea (*P*. *sativum*) that serve as representatives to examine KAI2 sub-functionalization in legumes. The identification of both *PsKAI2A* and *PsKAI2B* genes corroborates the recent finding that the *KAI2* gene duplication event occurred in Papilionoidaea before the diversification of legumes (*38*). The similarities in gene expression patterns are found between pea and *Lotus* with globally higher expression of *KAI2A* in comparison to *KAI2B* and higher expression *KAI2B* versus *KAI2A* in roots, depending on whether the plants are grown in pots or on Petri dishes. In the characterization of the PsKAI2 TILLING mutant lines, the effects on branching and root hair phenotypes were not significant. Thus, future studies of pea *Pskai2a/b* mutants will further delineate the distinct physiological functions, in particularly their symbiotic relationship with AM fungi and the differential expression patterns in the roots. Interestingly, both pea single mutants can induce the expression of *PsDLK2* in response to (–)-GR24, which is in agreement with the observation of (–)-GR24 perception by *L. japonicus* KAI2B (*38*) and substantiate the hindering effect of W158 in (–)-GR24 perception that is absent from pea PsKAI2s. Yet our DSF analyses suggest that PsKAI2B differs from its ortholog from *L. japonicus*, which is not destabilized by (–)-GR24 due to a rare phenylalanine to tryptophan exchange at position 158 at the binding pocket (*38*). The arabidopsis complementation experiments with pea KAI2s verified the distinct functionalities of PsKAI2A and PsKAI2B, wherein PsKAI2A most closely resembles AtKAI2 complementation in hypocotyl elongation assays indicating sensitivity to the endogenous KL, while PsKAI2B and not PsKAI2A is able to rescue the sensitivity of the *htl-3* and *kai2-2* mutants to (–)-GR24.

The occurrences of molecular coevolution of ligands and their specialized receptors have been previously demonstrated for phytohormones such as SL (*55*), ABA (*56*), GA (*57*), and more recently, karrikins (*35, 38*). Even though the exact identity of KL remains to be revealed, it is thought that the ligands likely share a common chemical composition to SLs. As such, it has been recently shown that one enantiomer of the artificial SL analogue, *rac*-GR24, can function by binding KAI2 in Arabidopsis (*9, 39, 49*). Here we show that PsKAI2B can form stronger interactions with the enantiomer (–)-GR24, compared to PsKAI2A. Moreover, we found that while both KAI2s are active hydrolases, they have distinct binding affinity and stereoselectivity towards GR24 stereoisomers. These findings indicate that sub-functionalization of KAI2s via substitutions in only few amino acids can greatly alter ligand affinity, binding, enzymatic activity, and probably signaling with downstream partners (*34, 38*).

To better elucidate PsKAI2A/B molecular divergence and their dual receptor-enzyme function, we carried out extensive biochemical and structural studies. KAI2/D14 crystal structures have greatly impacted our understanding of their receptor ligand-binding pockets and their ability to not only accommodate, but also hydrolyze certain ligands (*9, 21, 32, 34, 40–42*). The crystal structure of legume PsKAI2B together with the PsKAI2A homology model reported here, further substantiates the structural basis of this differential ligand selectivity. We identified conserved key amino acid changes that alter the shape of the pocket and confer altered ligand specificities. These atomic structures of legume KAI2 enabled us to analyze the distinction between key residues L160/S190/M218 in PsKAI2A and the corresponding residues M160/L190/L218 in PsKAI2B. Further swap experiments between residues 160 and 190 confirmed that these residues are necessary for the sensitivity of PsKAI2B for (–)-GR24, but not sufficient to bring PsKAI2A to similar sensitivities. These findings support recent *in planta* studies that demonstrate that residues 160 and 190 are required for differential ligand specificity between *Lotus* KAI2A and KAI2B (*38*). Furthermore, the residue in position 190 was also identified in the parasitic plant *Striga hermonthica* as being involved in forming differential specificity pockets between the highly variable and functionally distinct HTLs (*31, 32*). While the changes in positions 160 and 190 directly reshape the pocket morphology, the variant in position 218 is located in the center of the D-loop that has been suggested in downstream protein-protein interactions (*52, 53*). Therefore, the conserved substitution of KAI2A and KAI2B in M218 to L218 respectively across legumes may also contribute to downstream interaction(s). Based on the analogy with the D14-MAX2 perception mechanism, the KAI2 receptor is likely to adopt different conformational states upon ligand binding and cleavage. As such, the identification of unique residue variations in the lid (between KAI2A and KAI2B, respectively in positions 129 and 147) reported here, infer a sub-functionalization in the receptor regions that are likely to be involved in MAX2 and/or SMAX1 and/or SMXL2 downstream interactions. Therefore, it remains to be further elucidated whether these distinctive residues play a role in fine tuning the formation of the protein complex with MAX2-SMAX1/SMXL2.

The crystal structure of ligand-bound PsKAI2B provides a unique mechanistic view of perception and cleavage by KAI2s. Based on the crystallization conditions and following a detailed investigation of the electron density, we were able to overrule common chemicals and place the (–)-GR24 D-OH ring with higher relative fitting values than other components. The absence of positive electron density peaks corresponding to the intact (–)-GR24, and thus the presence of only the D-OH ring, raise questions of whether the S95-D-OH adduct recapitulates a pre- or post-cleavage intermediate state of (–)-GR24. The possibility that the trapped molecule represents a post cleavage state is intriguing and may provide a new intermediate state where S95 is covalently linked to the cleavage product. As such, the S95-D-OH adduct suggested here could explain the single turnover cycle that was observed for KAI2s in this study. While early structural study of D14 hydrolysis also positioned the catalytic serine with a covalent adduct, (*21*) other studies of the single turnover activity of D14 suggest that a covalent intermediate is in fact formed between the catalytic histidine and serine (*53*). The chemical similarity of the D-OH butenolide ring of karrikin and GR24 suggests that the KL signal may share a parallel structure and perhaps is biochemically processed via multiple steps and intermediate adducts. Therefore, the significance of this finding may also shed light on SL perception and cleavage by D14, which are still elusive. While the MS data corroborate a mass shift corresponding to an intermediate covalently bound to KAI2s, the adduct was detected more significantly on the catalytic histidine rather than on the serine. This data is in agreement with the expected transient nature of the serine nucleophilic attack, and the more stable adduct that can be formed on the catalytic histidine. Collectively, the crystal structure of PsKAI2B bound to enantiomeric SL synthetic analog and the MS data reveal, for the first time, the mode of action of KAI2 not only as receptors but also as *bona fide* enzymes. Beyond the importance of illuminating the stereoselectivity of ligand perception diverged KAI2 receptors in KAR/KL signaling pathways, our data strongly suggest the through the evolution of KAI2 enzymes, specific structural and functional adaptation diverged to enable more extended sensitivities to KAR/KL and SL and SL-like molecules by KAI2B.

Here, we elucidate the molecular basis for functional divergence of KAI2 receptors, focusing on a pea as model legume. Because of their ability to fix atmospheric nitrogen through plant–rhizobium symbiosis, legume crops such as pea or fava bean are attracting increasing attention for their agroecological potential. Thus, better understanding of KAR/KL perception and signaling in these key crops may have far-reaching impacts on agro-systems and food security.

## Methods

### Protein sequence alignment and phylogenetic tree analyses

Representative KAI2 sequences of 41 amino acid sequences were downloaded from Phytozome and specific genome databases as shown in **Fig. S1**. Alignment was performed in MEGA X (*58*) using the MUSCLE multiple sequence alignment algorithm (*59*). Sequence alignment graphics were generated using CLC Genomics Workbench v12. The evolutionary history was inferred by using the Maximum Likelihood method and JTT matrix-based model (*60*). Initial tree(s) for the heuristic search were obtained automatically by applying Neighbor-Join and BioNJ algorithms to a matrix of pairwise distances estimated using the JTT model, and then selecting the topology with superior log likelihood value. The percentage of trees in which the associated taxa clustered together is shown next to the branches (*61*). Tree is drawn to scale, with branch lengths measured in the number of substitutions per site. Analysis involved 41 amino acid sequences with a total of 327 positions in the final dataset. Evolutionary analyses were conducted in MEGA X (*58*).

### RT-PCR analyses

For PsKAI2A splicing variant detection, PCR reactions were performed using 1 µl of cDNA or genomic DNA sample in a final reaction mixture (20 µl) containing 2 µl of 10 x PCR buffer (ThermoFisher Scientific), 0.25 µL of 25 mM dNTPs, 0.25 µl of each primer at 10µM, and 1 unit of Dream Taq DNA polymerase (ThermoFisher Scientific). Primer sequences are indicated in **Table S3**. PCR was performed in the following conditions: 94 ◦C/5 min, 94 ◦C/30 s, 58 ◦C/30 s, 72 ◦C/1 min for 30 cycles. Half of each PCR product was loaded onto Ethidium bromide stained 1% agarose gels in TAE buffer, stained with ethidium bromide, and photographed with Molecular Imager^®^ Gel Doc™ XR System (BioRad)

### Constructs and generation of transgenic lines

The expression vectors for transgenic Arabidopsis were constructed by MultiSite Gateway Three-Fragment Vector Construction kit (Invitrogen). *AtKAI2* and *PsKAI2A.2* constructs were tagged with 6xHA epitope tag, mCitrine protein or GUS protein at their C-terminus. Lines were resistant to hygromycin. The *AtKAI2* native promoter (0.7 kb) was cloned into the pDONR-P4P1R vector, using Gateway recombination (Invitrogen) as described in (*62*).The 6xHA with linker and mCitrine tags were cloned into pDONR-P2RP3 (Invitrogen) as described in de Saint Germain et al. (*48*). *PsKAI2A.1*, *PsKAI2A.2* and *PsKAI2B* CDS were PCR amplified from *Pisum* cv. Térèse cDNA with the primers specified in **Table S3**.and then recombined into the pDONR221 vector (Invitrogen). The suitable combination of *AtKAI2* native promoter, *AtKAI2*, *PsKAI2A.1*, *PsKAI2A.2* or *PsKAI2B* and 6XHA,mCitrine or GUS was cloned into the pH7m34GW final destination vectors by using the three fragment recombination system (*63*) and were thusly named proAtKAI2:AtKAI2-6xHA, proAtKAI2:AtKAI2-mcitrine, proAtKAI2:AtKAI2-GUS, proAtKAI2:PsKAI2A.1-6xHA, proAtKAI2:PsKAI2A.2-mcitrine, proAtKAI2:PsKAI2A.2-GUS, proAtKAI2:PsKAI2B-6XHA, proAtKAI2:PsKAI2B-GUS and proAtKAI2:PsKAI2B-mcitrine.. Transformation of Arabidopsis *htl-3* or *kai2-2* mutant was performed according to the conventional floral dipping method (*64*), with Agrobacterium strain GV3101. For each construct, only a few independent T1 lines were isolated, and all lines were selected in T2. Phenotypic analysis shown in **Fig. 1E** was performed on the T3 homozygous lines.

### Protein extraction and immunoblotting

Total protein extract was prepared from 8 to 10, 10 day-old Arabidopsis seedling in Laemmli buffer and boiled for 5 min. Total protein were separated by 10% SDS-PAGE and transferred onto polyvinylidene difluoride membrane (Bio-Rad) probed with anti-HA primary antibody (1:10000; SIGMA H9658-100UL Lot#128M4789V) and then anti-mouse-IgG-HRP secondary antibody (1:10000; SIGMA A9044-2ML-100UL Lot#029M4799V) or with anti-GFP primary antibody (1:10000; CHROMTEK 3H9-100 Lot#60706001AB) and then anti-rat-IgG-HRP secondary antibody (1:10000; SIGMA A9037-1ML Lot#SLCF6775). Ponceau staining was used as a loading control.

### Identification of Pskai2a and Pskai2b Targeting-Induced Local Lesions IN Genomes (TILLING) mutants

The mutagenized population in the pea cultivar (cv.) Caméor was used as a TILLING resource. For obtaining mutants in *PsKAI2A*, TILLING analysis was performed on 5000 families within one amplicon of 1068 bp using nested primers (N1, N2) with PsKAI2A_N1F primer and PsKAI2A_N1R primer PsKAI2A_N2Ftag primer and PsKAI2A_N2Rtag primer Primers are indicated in **Table S3**. The enzymatic mutation detection technique based on the mismatch specific endonuclease ENDO1 was used. For *PsKAI2B*, the mutation detection system by sequencing and described in (*65*) was used. Two amplicons of 381 and 401 bp were screened in 2500 families. Primers are indicated in **Table S3**. Prediction of the amino acid changes that affect protein function was made using the SIFT program (sift.jcvi.org/).

M3 and M4 seeds from lines carrying mutations in the *PsKAI2A* and *PsKAI2B* genes were genotyped for homozygous mutant plants; these plants were backcrossed once (alleles *Pskai2a-4, Pskai2b-1, Pskai2b-2*) to three or four times (alleles *Pskai2a-2, Pskai2a-6, Pskai2b-3*) to the cv. Caméor. BC1-F3 and M5 single mutant plants were crossed for obtaining the *Pskai2a-6 Pskai2b-3* double mutant.

### Plant material and growth conditions

For branching quantification, *Pisum sativum* plants were grown in glasshouse (23°C day/ 15°C night) under a 16-h photoperiod (the natural daylength was extended or supplemented during the day when necessary using sodium lamps) in pots filled with clay pellets, peat, and soil (1:1:1) supplied regularly with nutrient solution. Nodes were numbered acropetally from the first scale leaf as node 1.

### Root hair assay

*Pisum sativum Cameor* wild-type, *Pskai2a-3*, *Pskai2b-6*, *Pskai2a-3 Pskai2a-6* seeds were surface sterilized with 1% NaClO, washed 5 times, and incubated for 2 hours in sterile water. Imbibed seeds were germinated on ½ MS, pH 5.8 containing 1% agar at 4°C for 3 days in the dark. Seedlings were grown in axenic conditions on 12×12 cm square Petri dishes at 24°C with 16-h-light/8-h-dark cycles. To assess root hair length, images of the primary root tips of 10-day old seedlings were taken with a Zeiss Discovery V8 microscope equipped with a Zeiss Axiocam 503 camera. Root hair length was measured for a minimum of 8 roots per genotype for 8 different root hairs per root, between 10 and 20 mm from the root tip using Fiji as described (*66*). For root-hair length measurements a Welch t-test, p-value < 0.05 and for RT-qPCR analysis a Kruskal-Wallis Test with post-hoc Student’s t-test, p<0.05 were performed using R statistical environment (https://www.r-project.org/). For the Kruskal-Wallis Test the R-package agricolae (https://CRAN.R-project.org/package=agricolae) was used.

### Treatment for analysis of transcript accumulation

For treatments with KARs and GR24 enantiomers, 10-day old seedlings grown on Petri dishes as described above, were placed with their roots into 50 ml amber Falcon tubes filled with ½ MS solution for 24 h to allow the seedlings to adapt to the new growth system. For the treatment the growth media was exchanged with ½ MS solution containing 3μM Karrikin_1_, Karrikin_2_, (www.olchemim.cz), (+)-GR24 or (-)-GR24 (www.strigolab.eu) and seedlings were incubated with their shoots in the light for 4 hours.

### Analysis of transcript accumulation by RT-qPCR

For analysis of transcript levels by RT-qPCR presented in figure 1, total RNA was isolated from 28 days old plant for flower and flower bud and from 10 days old plants for all other tissues, using TRIZOL reagent (Invitrogen) following the manufacturer’s protocol. DNase treatment was performed to remove DNA using the Qiagen RNase-Free DNase Set (79254) and the RNeasy Mini Kit (74904) and eluted in 50 mL of RNase-free water. RNA was quantified using NanoDrop 1000 and migrated on gels to check RNA non-degradation. Total cDNA was synthesized from 2 mg of total RNA using 50 units of RevertAid H Moloney murine leukemia virus reverse transcriptase in 30 µL following the manufacturer’s instructions with poly(T)18 primer. cDNA was diluted 10 times before subsequent analysis. Quantitative reverse transcription-PCR analyses were adapted from (*67*). They were performed using SsoAdvancedTM Universal SYBR® Green SuperMix (Biorad). Cycling conditions for amplification were 95°C for 10 min, 50 cycles of 95°C for 5 s, 62°C for 5 s, and 72°C for 15 s, followed by 0.1°C s–1 ramping up to 95°C for fusion curve characterization. Two biological repeats were analyzed in duplicate. To calculate relative transcript levels, the comparative cycle method based on non-equal efficiencies was used (*68*). Transcript levels for the different genes were expressed relative to the expression of the *PsACTIN* gene. Primers are indicated in **Table S3**.

For analysis of transcript levels by RT-qPCR presented in **Fig. 2**, plant tissue was rapidly shock frozen in liquid nitrogen and ground to a fine powder with a mortar and pestle. RNA was extracted using the Spectrum Plant Total RNA Kit (www.sigmaaldrich.com). The RNA was treated with Invitrogen DNAse I amp. grade (www.invitrogen.com) and tested for purity by PCR. cDNA synthesis was performed with 1 µg RNA using the iScript cDNA Synthesis kit (www.Biorad.com). cDNA was diluted in water in a 1:10 ratio and 1µl was used for RT-PCR was performed with an iCycler (Biorad, www.bio-rad.com/) using a Green MasterMix (Jena Bioscience, highROX, 2x conc.). Thermal cycler conditions were: 95°C 2 min, 45 cycles of 95°C 30 sec, 60°C 30sec and 72°C 20 sec followed by dissociation curve analysis. Expression levels were calculated according to the ΔΔCt method (*69*). For each genotype and treatment three to four biological replicates were monitored and each sample was represented by two technical replicates. Transcript levels for the different genes were expressed relative to the expression of the *PsTUB* gene, Accession:X54844, (*70*). Primers are indicated in **Table S3**.

### Hypocotyl elongation assays

Arabidopsis seeds were surface sterilized by consecutive treatments of 5 min 70% (v/v) ethanol with 0.05% (w/v) sodium dodecyl sulfate (SDS) and 5 min 95% (v/v) ethanol. Then seeds were sown on half-strength Murashige and Skoog (½ MS) media (Duchefa Biochemie) containing 1% agar, supplemented with 1 μM (–)-GR24 or with 0.01 % DMSO (control). Seeds were stratified at 4 °C (2 days in dark) then transferred to the growth chamber at 22 °C, under 20-30 µE /m^2^/sec of white light in long day conditions (16 hr light/ 8 hr dark). Seedlings were photographed and hypocotyl lengths were quantified using ImageJ (*71*). 2 plates of 10-12 seeds were sown for each genotype x treatment. Using Student t-tests, no statistically significantly different means were detected between plates. The data from the 20-24 seedlings were then used for a one-way ANOVA.

### Chemicals

Enantiopure GR24 isomers were obtained as described in de Saint Germain et al. (*48*) or purchased from StrigoLab. Karrikin_1_ and Karrikin_2_ were purchased from Olchemim. Profluorescent probes (GC240, GC486) were obtained as described in de Saint Germain et al. (*48, 72*).

### Protein preparation and purification

PsKAI2A.2, PsKAI2B, and all described mutants were independently cloned and expressed as a 6× His-SUMO fusion proteins from the expression vector pAL (Addgene). These were cloned utilizing primers in **Table S3**. BL21 (DE3) cells transformed with the expression plasmid were grown in LB broth at 16 °C to an OD_600_ of ∼0.8 and induced with 0.2 mM IPTG for 16 h. Cells were harvested, re-suspended and lysed in extract buffer (50 mM Tris, pH 8.0, 200 mM NaCl, 5 mM imidazole, 4% Glycerol). All His-SUMO-PsKAI2s were isolated from soluble cell lysate by Ni-NTA resin. The His-SUMO-PsKAI2 was eluted with 250 mM imidazole and subjected to anion-exchange. The eluted protein was than cleaved with TEV (tobacco etch virus) protease overnight at 4 °C. The cleaved His-SUMO tag was removed by passing through a Nickel Sepharose and PsKAI2 was further purified by chromatography through a Superdex-200 gel filtration column in 20 mM HEPES, pH 7.2, 150 mM NaCl, 5 mM DTT, 1% Glycerol. All proteins were concentrated by ultrafiltration to 3–10 mg/mL^−1^. RMS3, AtD14, AtKAI2 were expressed in bacteria with TEV cleavable GST tag, purified and used as described in de Saint Germain et al. (*48*).

### Enzymatic hydrolysis of GR24 isomers by purified proteins

Ligands (10 µM) were incubated without and with purified proteins (5 µM) for 150 min at 25 °C in PBS (0.1 mL, pH 6.8) in presence of (±)-1-indanol (100 µM) as the internal standard. The solutions were acidified to pH 1 with 10% trifluoroacetic acid in CH_3_CN (v/v) (2 µL) to quench the reaction and centrifuged (12 min, 12,000 tr/min). Thereafter the samples were subjected to RP-UPLC-MS analyses using Ultra Performance Liquid Chromatography system equipped with a PDA and a Triple Quadrupole mass spectrometer Detector (Acquity UPLC-TQD, Waters, USA). RP-UPLC (HSS C_18_ column, 1.8 μm, 2.1 mm × 50 mm) with 0.1% formic acid in CH_3_CN and 0.1% formic acid in water (aq. FA, 0.1%, v/v, pH 2.8) as eluents [10% CH_3_CN, followed by linear gradient from 10 to 100% of CH_3_CN (4 min)] was carried out at a flow rate of 0.6 mL/min. The detection was performed by PDA using the TQD mass spectrometer operated in Electrospray ionization positive mode at 3.2 kV capillary voltage. The cone voltage and collision energy were optimized to maximize the signal and were respectively 20 V for cone voltage and 12 eV for collision energy and the collision gas used was argon at a pressure maintained near 4.5.10^-3^ mBar.

### Enzymatic assay with pro-fluorescent probes

Enzymatic assay and analysis have been carried out as described in de Saint Germain et al. (*48*), using a TriStar LB 941 Multimode Microplate Reader from Berthold Technologies. The experiments were repeated three times.

### Protein melting temperatures

Differential Scanning Fluorimetry (DSF) experiments were performed on a CFX96 TouchTM Real-Time PCR Detection System (Bio-Rad Laboratories, Inc., Hercules, California, USA) using excitation and emission wavelengths of 490 and 575 nm, respectively. Sypro Orange (λ_ex_/λ_em_ : 470/570 nm; Life Technologies Co., Carlsbad, California, USA) was used as the reporter dye. Samples were heat-denatured using a linear 25 to 95 °C gradient at a rate of 1.3 °C per minute after incubation at 25 °C for 30 min in the absence of light. The denaturation curve was obtained using CFX manager™ software. Final reaction mixtures were prepared in triplicate in 96-well white microplates, and each reaction was carried out in 20 μL scale in Phosphate buffer saline (PBS) (100 mM Phosphate, pH 6.8, 150 mM NaCl) containing 6 μg protein (such that final reactions contained 10 μM protein), 0-1000 μM ligand (as shown on the **Fig. 4c-f** **and Fig. S8b-e**), 4% (v/v) DMSO, and 0.008 μL Sypro Orange. Plates were incubated in darkness for 30 minutes before analysis. In the control reaction, DMSO was added instead of ligand. All experiments were repeated three times.

### Intrinsic tryptophan fluorescence assays and kinetics

Intrinsic tryptophan fluorescence assays and determination of the dissociation constant K_D_ has been performed as described in de Saint Germain et al. (*48*), using the Spark® Multimode Microplate Reader from Tecan.

### Crystallization, data collection and structure determination

The crystals of PsKAI2B were grown at 25 °C by the hanging-drop vapor diffusion method with 1.0 μL purified protein sample mixed with an equal volume of reservoir solution containing 0.1 M HEPES pH 7.5, 2.75% PEG 2000, 2.75% v/v PEG 3350, 2.75% v/v PEG 4000, 2.75% v/v PEG-ME 5000. The crystals of PsKAI2B in complex with (–)-GR24 were grown at 25 °C by the hanging-drop vapor diffusion method with 1.0 μL purified protein complex (preincubated with 1 mM (–)-GR24, StrigoLab) and mixed with an equal volume of reservoir solution containing 0.1 M HEPES pH 7.5, 2.75% PEG 2000, 2.75% v/v PEG 3350, 2.75% V/V PEG 4000, 2.75% v/v PEG-ME 5000, 1 mM (–)-GR24. Crystals of maximum size were obtained and harvested after 2 weeks from the reservoir solution with additional 20% MPD serving as cryoprotectant. X-ray diffraction data was integrated and scaled with HKL2000 package (*73*). PsKAI2s crystal structures were determined by molecular replacement using the AtKAI2 model (PDB: 5Z9H) (*74*) as the search model. All structural models were manually built, refined, and rebuilt with PHENIX (*75*) and COOT (*76*).

### Structural biology modelling and analyses

Model structure illustrations were made by PyMOL (*77*). PsKAI2A model structure was generated using iTASSER (*78*). Ligand identification, ligand-binding pocket analyses, and computing solvent accessible surface values analyses were carried out using Phenix LigandFit (*75*), CASTp software (*79*), and AutoDock Vina (*80*), respectively. LigPlot+ program (*81*) was used for 2-D representation of protein-ligand interactions from standard PDB data format.

### Direct electrospray ionization – mass spectrometry of PsKAI2 proteins (ESI)-MS under denaturing conditions

Mass spectrometry measurements were performed with an electrospray Q-TOF mass spectrometer (Waters) equipped with the Nanomate device (Advion, Inc.). The HD_A_384 chip (5 μm I.D. nozzle chip, flow rate range 100−500 nL/min) was calibrated before use. For ESI−MS measurements, the Q-TOF instrument was operated in RF quadrupole mode with the TOF data being collected between m/z 400−2990. Collision energy was set to 10 eV and argon was used as the collision gas. PsKAI2 proteins (50 µM) in 50 mM ammonium acetate (pH 6.8) in presence or without (-)_GR24 (500 µM) were incubated for 10 min at room temperature before denaturation in 50% acetonitrile and 1% formic acid. The solutions were directly injected for Mass spectra acquisition or digested before LC-MS/MS analyses. Mass Lynx version 4.1 (Waters) and Peakview version 2.2 (Sciex) software were used for acquisition and data processing, respectively. Deconvolution of multiply charged ions was performed by applying the MaxEnt algorithm (Sciex). The average protein masses were annotated in the spectra and the estimated mass accuracy was ± 2 Da. External calibration was performed with NaI clusters (2 μg/μL, isopropanol/H_2_O 50/50, Waters) in the acquisition m/z mass range.

## Acknowledgements

N.S. is supported by NSF-CAREER (Award #2047396) and NSF-EAGER (Award #2028283). This work is also supported by the CHARM3AT Labex program (ANR-11-LABX-39) to F.-D.B.; by AgreenSkills from the European Union in the framework of the Marie-Curie FP7 COFUND People Programme and a fellowship from Saclay Plant Sciences (ANR-17-EUR-0007) to A.d.S.G.; and by the Emmy Noether program (GU1423/1-1) of the Deutsche Forschungsgemeinschaft (DFG) to C.G.. Furthermore, this work was supported by the Institut Jean-Pierre Bourgin’s Plant Observatory technological platforms. We thank the beamline staff at the Advanced Light Source (U.S. DOE Office of Science User Facility under Contract No. DE-AC02-05CH11231, is supported in part by the ALS-ENABLE program funded by the National Institutes of Health, National Institute of General Medical Sciences, grant P30 GM124169-01). We thank the facilities and expertise of the I2BC proteomic platform (Proteomic-Gif, SICaPS) supported by IBiSA, Ile de France Region, Plan Cancer, CNRS and Paris-Sud University

## Author Contributions

AM.G., S.T., F.-D.B., C.R., C.G., A.dS.G., and N.S. conceived and designed the experiments. N.S., A.dS.G., and AM.G. conducted the protein purification, biochemical and crystallization experiments. J.-P.P. characterized pea Tilling mutants. S.T. performed pea RT-qPCR and root hair assay. P.LB. obtained Arabidopsis complementation lines. A.B., M.D. and C.LS. generated pea TILLING mutant collection. D.C. performed the mass experiments. N.S. and AM.G. determined and analyzed crystal structures and conducted in silico studies. S. T. performed pea root hair assays and gene expression analysis in **Fig. 2**. AM.G., A.dS.G., and N.S. wrote the manuscript with help from S.T., F.-D.B., C.R. and C.G..

## Data and materials availability

The atomic coordinates of apo and ligand-bound forms of PsKAI2 structures has been deposited in the Protein Data Bank with accession codes 7K2Z and 7K38, respectively. All relevant data are available from corresponding authors upon request.

