## Supplementary Materials for "Structural and Functional Analyses Explain Pea KAI2 Receptor Diversity and Reveal Stereoselective Catalysis During Signal Perception"

**Fig. S1.** Proteins Sequence analysis of KAI2/D14 family.

**Fig. S2.** Alternative splicing of the *PsKAI2A* transcript.

**Fig. S3.** Multiple sequence alignment and conservation analysis of representative legume and non-legume KAI2s.

**Fig. S4.** TILLING mutant residues position on KAI2 structures.

**Fig. S5.** Branching and root hair phenotype *Pskai2* mutants.

**Fig. S6.** Hypocotyl elongation in *kai2-2* mutant and KAI2s protein expression in complementation assay.

**Fig. S7**. Purification of PsKAI2s.

**Fig. S8.** Biochemical and structural analysis of the interaction between PsKAI2 and 2' isomers ligands.

**Fig. S9.** Intrinsic tryptophan fluorescence of PsKAI2s, AtKAI2 and RMS3 proteins in the presence of SL analogs.

**Fig. S10.** Structural divergence analysis of legume KAI2A and KAI2B.

**Fig. S11.** Biochemical and structural analysis of swap mutant PsKAI2A/B and their interaction with (–)-GR24.

**Fig. S12.** Structural interrogation of the ligand bound PsKAI2B crystal structure.

**Fig. S13.** Mass spectrometry characterization of covalent PsKAI2-ligand complexes.

**Table S1.** PsKAI2 TILLING mutants.

**Table S2.** Data collection, phasing and refinement statistics.

**Table S3.** Primer sequences used in study.

**a**

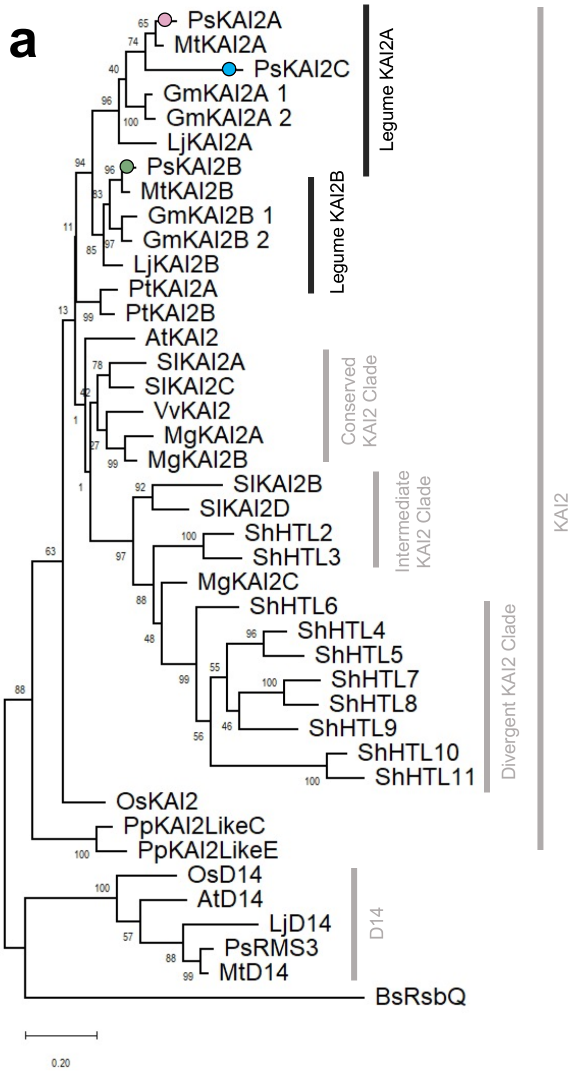

**b**

| | Species | Gene name | Sequence ID | Source |
|---|---|---|---|---|
| | Pisum sativum | KAI2A | Psat2g169960 | ANR GENOPEA |
| | Pisum sativum | KAI2B | Psat4g083040 | ANR GENOPEA |
| | Pisum sativum | KAI2C | Psat3g014200 | ANR GENOPEA |
| | Oryza sativa | KAI2 | LOC_Os03g32270.1 | Phytozome |
| | Arabidopsis thaliana | KAI2 | AT4G37470.1 | TAIR |
| | Populus trichocarpa | KAI2A | Potri.005G145000.1 | Phytozome |
| | Populus trichocarpa | KAI2B | Potri.007G052000.1 | Phytozome |
| | Lotus japonicus | KAI2A | Lj2g3v1931930.1 | LORE1 |
| | Lotus japonicus | KAI2B | Lj0g3v0117039.1 | LORE1 |
| | Solanum Lycopersicon | KAI2A | Solyc02g064770.2.1 | Phytozome |
| | Solanum Lycopersicon | KAI2B | Solyc02g064760.1.1 | Phytozome |
| | Solanum Lycopersicon | KAI2C | Solyc02g092770.2.1 | Phytozome |
| | Solanum Lycopersicon | KAI2D | Solyc02g092760.2.1 | Phytozome |
| | Vitis vinifera | KAI2 | GSVIVT01000162001 | Phytozome |
| | Mimulus guttatus | KAI2A | Migut.H00375.1 | Phytozome |
| | Mimulus guttatus | KAI2B | Migut.H00378.1 | Phytozome |
| | Mimulus guttatus | KAI2C | Migut.L00194.1 | Phytozome |
| KAI2 | Physcomitrella patens | KAI2-LikeC | 6ATX | RCSB |
| | Physcomitrella patens | KAI2-LikeE | 6AZB | RCSB |
| | Striga hermonthica | HTL2 | from Toh et al 2015 | Toh et al 2015[56] |
| | Striga hermonthica | HTL3 | from Toh et al 2015 | Toh et al 2015[56] |
| | Striga hermonthica | HTL4 | from Toh et al 2015 | Toh et al 2015[56] |
| | Striga hermonthica | HTL5 | from Toh et al 2015 | Toh et al 2015[56] |
| | Striga hermonthica | HTL6 | from Toh et al 2015 | Toh et al 2015[56] |
| | Striga hermonthica | HTL7 | from Toh et al 2015 | Toh et al 2015[56] |
| | Striga hermonthica | HTL8 | from Toh et al 2015 | Toh et al 2015[56] |
| | Striga hermonthica | HTL9 | from Toh et al 2015 | Toh et al 2015[56] |
| | Striga hermonthica | HTL10 | from Toh et al 2015 | Toh et al 2015[56] |
| | Striga hermonthica | HTL11 | from Toh et al 2015 | Toh et al 2015[56] |
| | Glycine max | KAI2A 1 | Glyma.01G191200.1.p | Phytozome |
| | Glycine max | KAI2A 2 | Glyma.11G051000.1.p | Phytozome |
| | Glycine max | KAI2B 1 | Glyma.17G164500.1.p | Phytozome |
| | Glycine max | KAI2B 2 | Glyma.05G102800.1.p | Phytozome |
| | Medicago truncatula | KAI2A | Medtr5g016150.1 | Phytozome |
| | Medicago truncatula | KAI2B | Medtr4g095310.1 | Phytozome |
| | Pisum sativum | RMS3 | Psat6g018320 | ANR GENOPEA |
| | Oryza sativa | D14 | 3W04 | RCSB |
| D14 | Arabidopsis thaliana | D14 | AT3G03990 | TAIR |
| | Lotus japonicus | D14 | Lj5g3v0310140.4 | LORE1 |
| | Medicago truncatula | D14 | Medtr1g018320.1 | Phytozome |
| RsbQ | Bacillus subtilis | RsbQ | WP_187001895 | NCBI |

**Fig. S1. Proteins Sequence analysis of KAI2/D14 family.** (**a**) Maximum likelihood phylogeny of 41 KAI2/D14-family proteins with bacterial RBSQ as outgroup. Node values represent percentage of trees in which the associated taxa clustered together. Vertical rectangles highlight distinct *KAI2/D14* family clades. Black circle indicates legume duplication event. Pink and green circles mark the position of *PsKAI2As* and *PsKAI2B* respectively. The tree is drawn to scale, with branch lengths measured in the number of substitutions per site. (**b**) Sequence identifiers and sources for each.

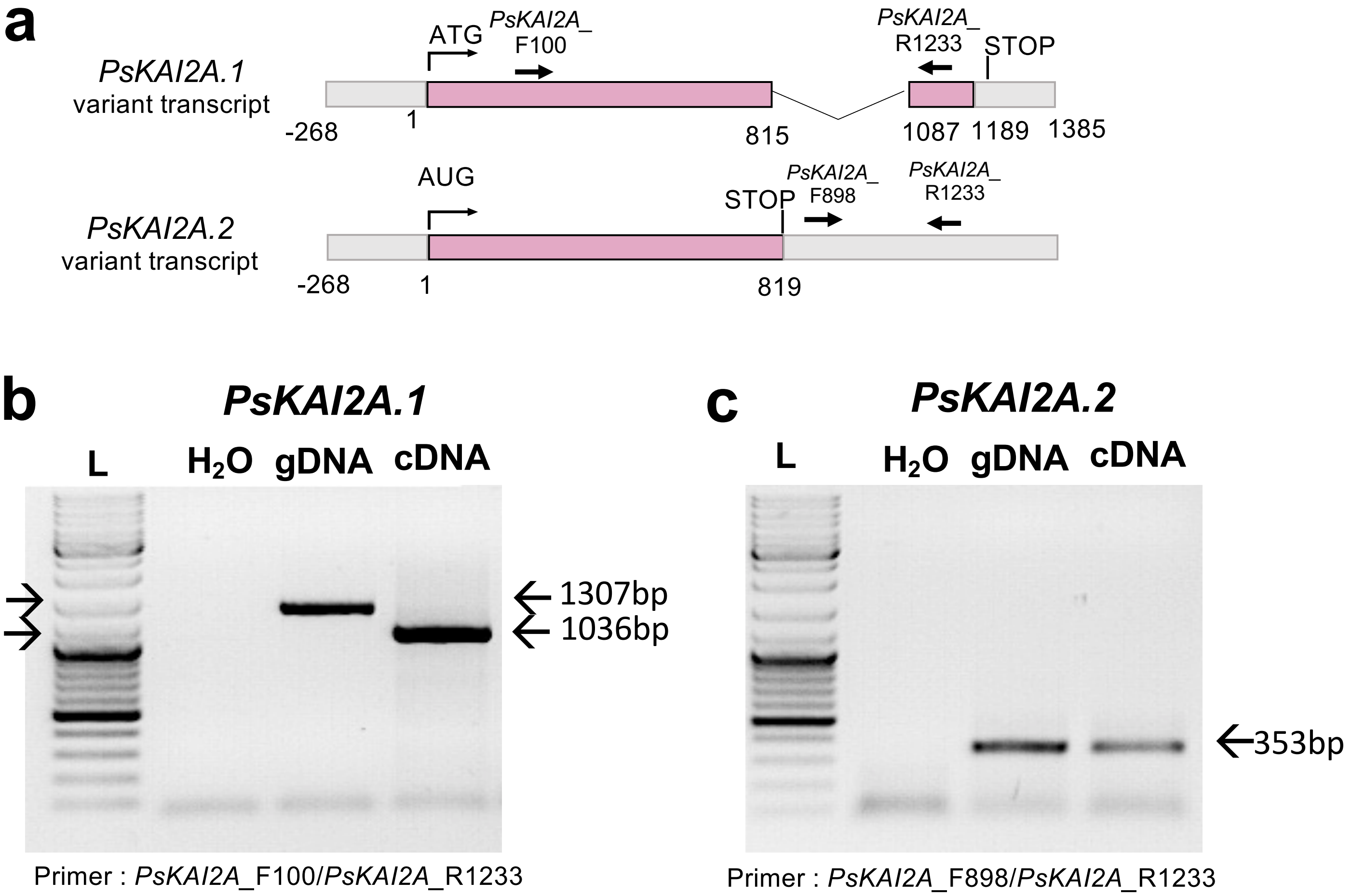

**Fig. S2. Alternative splicing of the *PsKAI2A* transcript.** (**a**) Schematic representation of the splicing events leading to *PsKAI2A.1* and *PsKAI2A.2* forms of mature transcripts. Exons are indicated as pink boxes. (**b-c**) Electrophoresis gel of PCR products obtained after amplification of the PsKAI2A coding sequence with primer specific to *PsKAI2A.1 f*orm (**b**) and primer specific to *PsKAI2A.2 f*orm (**c**) using genomic DNA (gDNA) or first-strand cDNA (cDNA) as template; H2O: negative control; Lane L: DNA ladder.

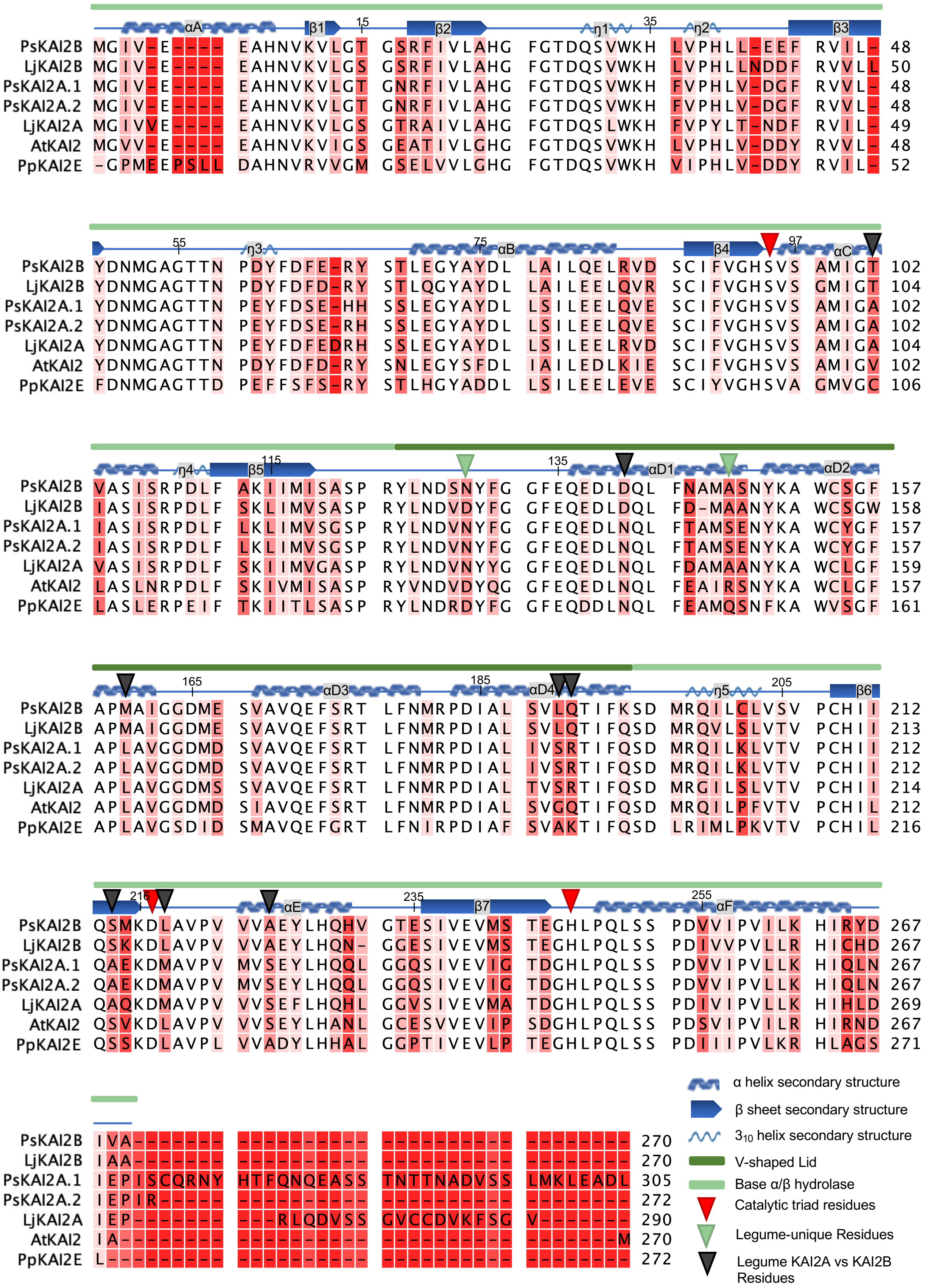

**Fig. S3. Multiple sequence alignment and conservation analysis of representative legume and non-legume KAI2s.** Multiple sequence alignment and conservation analysis of selected KAI2s. Amino acid alignment of 7 plant KAI2 proteins. Intensity of red behind residues shows degree of divergence. Numbers on residues refer to position in PsKAI2B sequence. PsKAI2B lid (forest green) and base (light green) domains are indicated above alignment. Secondary structure of PsKAI2B sequence is shown above sequence in blue as alpha helices (αA- αF, base; and αD1- αD4, lid), beta sheets (β1- β7), 310 helices (η1- η5) and non-secondary structure-containing loops. Red arrows indicate catalytic triad residues, green arrows indicate legume KAI2-unique residues as shown in **Fig. 6**, dark grey arrows indicate residues diverged between legume KAI2A and KAI2B as described in **Fig. 7** and **Fig. S10**.

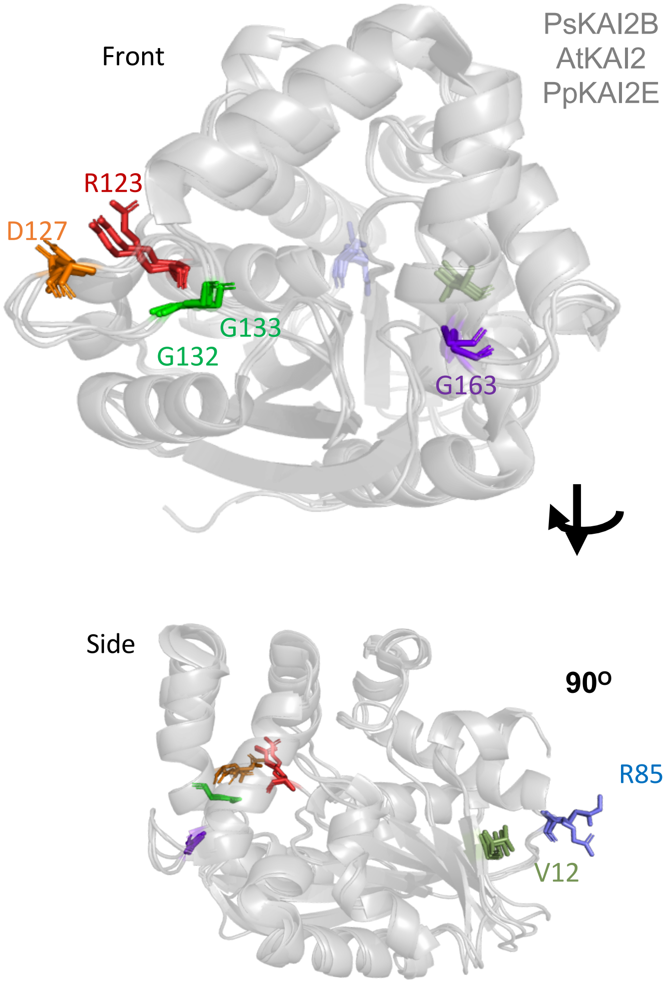

**Fig. S4. TILLING mutant residues position on KAI2 structures.** Superposition of PsKAI2B. AtKAI2, and PpKAI2E structures are shown in grey. The identified TILLING mutant residues are highlighted and labelled in rainbow colors using PsKAI2B as reference for amino acid sequence position.

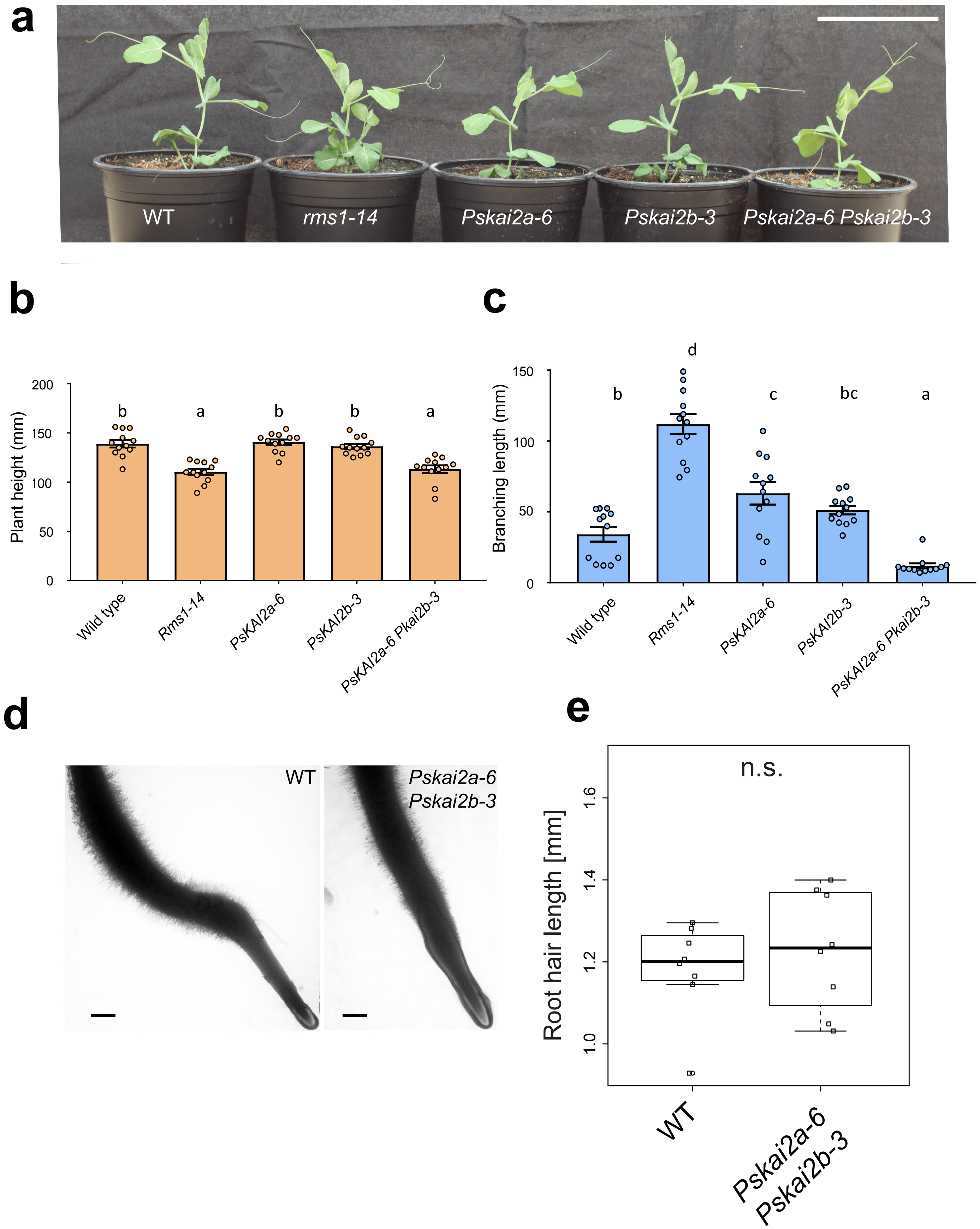

**Fig. S5. Branching and root hair phenotype *Pskai2* mutants.** (**a**) Phenotype and quantification of (**b**) plant height and (**c**) plant lateral length (mm) of 10-day-old *Pisum sativum* plant. Bar, 8.5 cm. Data are means ± SE (n=11-12). Statistical differences were determined using a one-way ANOVA with a Tukey multiple comparison of means post-hoc test, statistical differences of P<0.05 are represented by different letters. (**d**) Root hair length phenotype and quantification (**e**) of 10 day old WT and *Pskai2a-6 Pskai2b-3* seedlings. Bars, 2 mm. Data are means of measurements (n=8) of individual root (n=8-9), Statistical analysis Welch t-test, p-value < 0.05

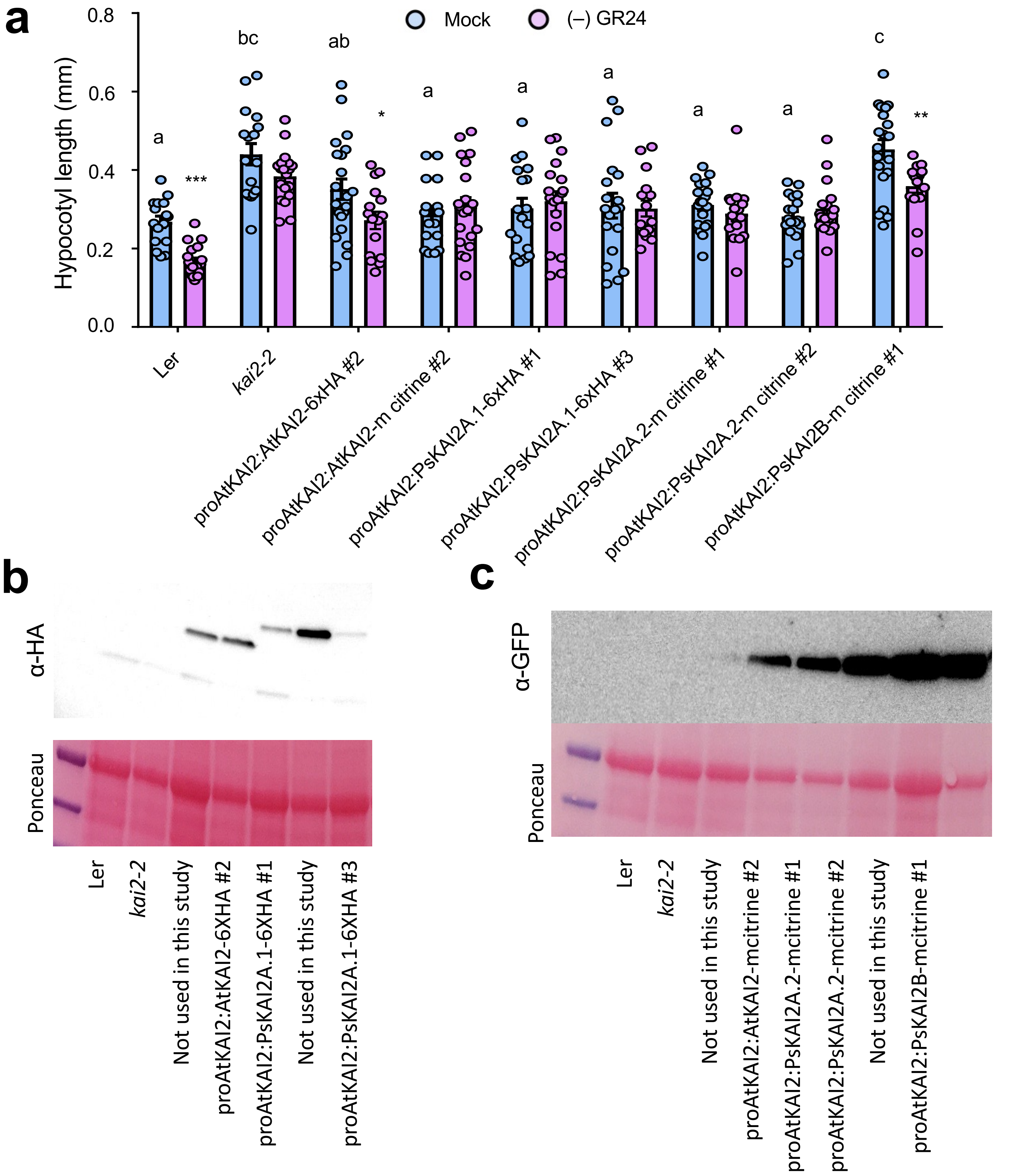

**Fig. S6. Hypocotyl elongation in *kai2-2* mutant and KAI2s protein expression in complementation assay. (a)** Hypocotyl length of 7-day-old seedlings grown under low light at 21 °C. Data are means ± SE (n = 20-24; 2 plates of 10-12 seedlings per plate). Light blue bars: Mock (DMSO), lavender bars: (–)-GR24 (1μM). Complementation assays using the *AtKAI2* promoter to express *AtKAI2* (control) or *PsKAI2* genes in the null *kai2-2* mutant background (Ler ecotype) as noted above the graph. Proteins were tagged with 6xHA epitope or mCitrine protein. For DMSO controls, statistical differences were determined using a one-way ANOVA with a Tukey multiple comparison of means post-hoc test, statistical differences of $P<0.05$ are represented by different letters. Means with asterisks indicate significant inhibition compared to mock-treated seedlings with *** corresponding to $p \leq 0.001$ and * to $p \leq 0.01$ , as measured by t-test. **(b-c)** AtKAI2 and PsKAI2 level analyzed by immunoblot using α-HA antibody **(b)** or α-GFP antibody **(c)** in Col-0 (wild type), *kai2-2* and transformed *kai2-2* plants, expressed under the control of *AtKAI2* promoter. Protein extracts from 10-d-old seedlings were separated by 10% SDS-PAGE. Ponceau staining is included for loading reference.

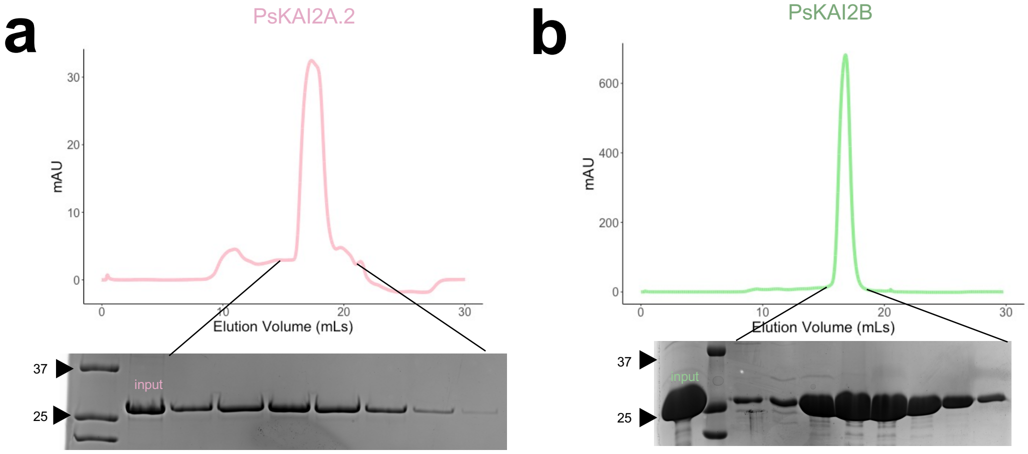

**Fig. S7. Purification of PsKAI2s.** Size exclusion peaks eluates for PsKAI2A.2 (**a**) and PsKAI2B (**b**). Proteins were resolved on SDS-PAGE gels and visualized via Coomassie stain. Molecular weight (MW) markers are labeled to show the relative size of proteins.

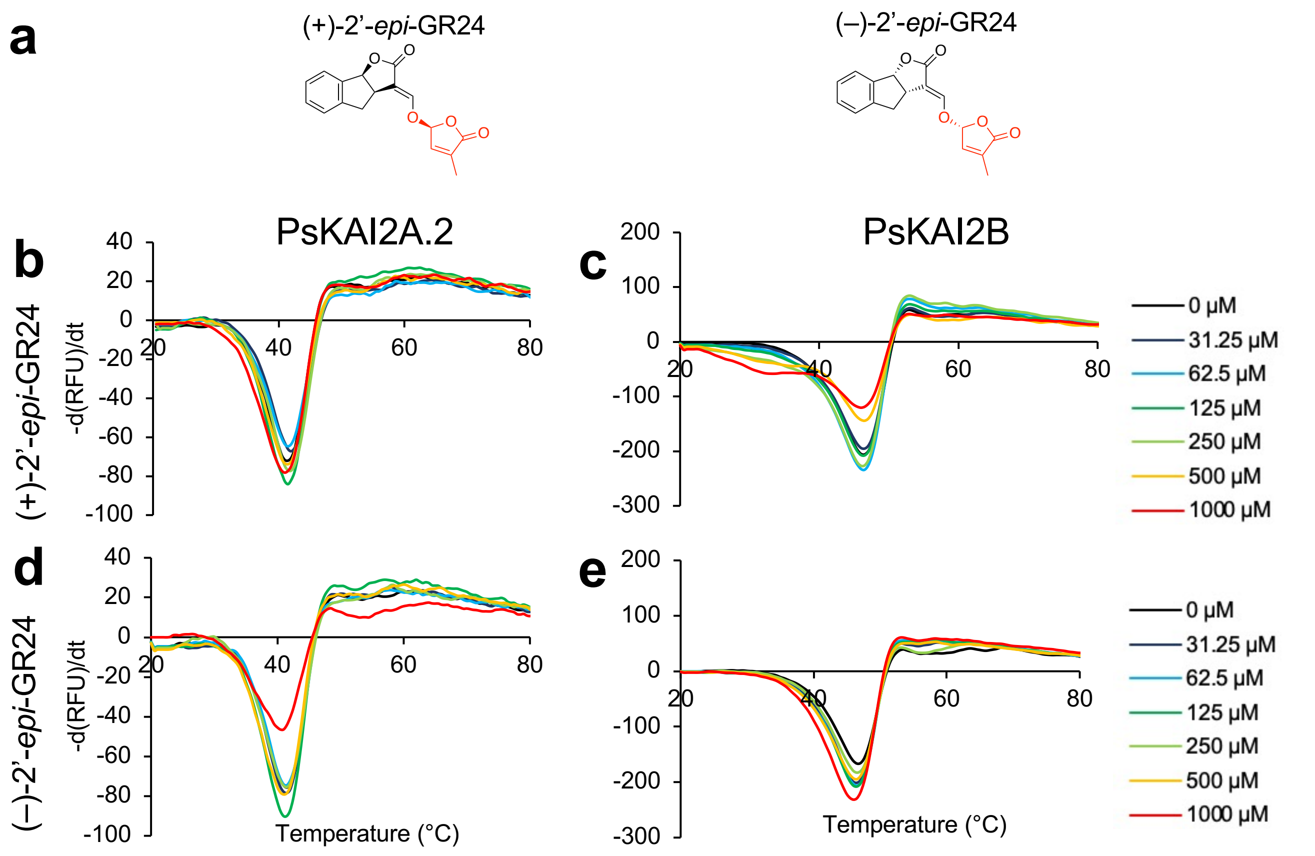

**Fig. S8. Biochemical analysis of the interaction between PsKAI2 proteins and the (+)-2’-*epi*-GR24 and the (-)-2’-*epi*-GR24 isomers ligands by DSF.** (**a**) Chemical structure of ligands used in DSF assay. The melting temperature curves of 10 µM PsKAI2A.2 (**b**, **d**) or PsKAI2B (**c**, **e**) with (+)-*2’-epi*-GR24 (b-c), or (–)-*2’-epi*-GR24 (**d-e**) at varying concentrations are shown as assessed by DSF. Each line represents the average protein melt curve for three technical replicates and the experiment was carried out twice.

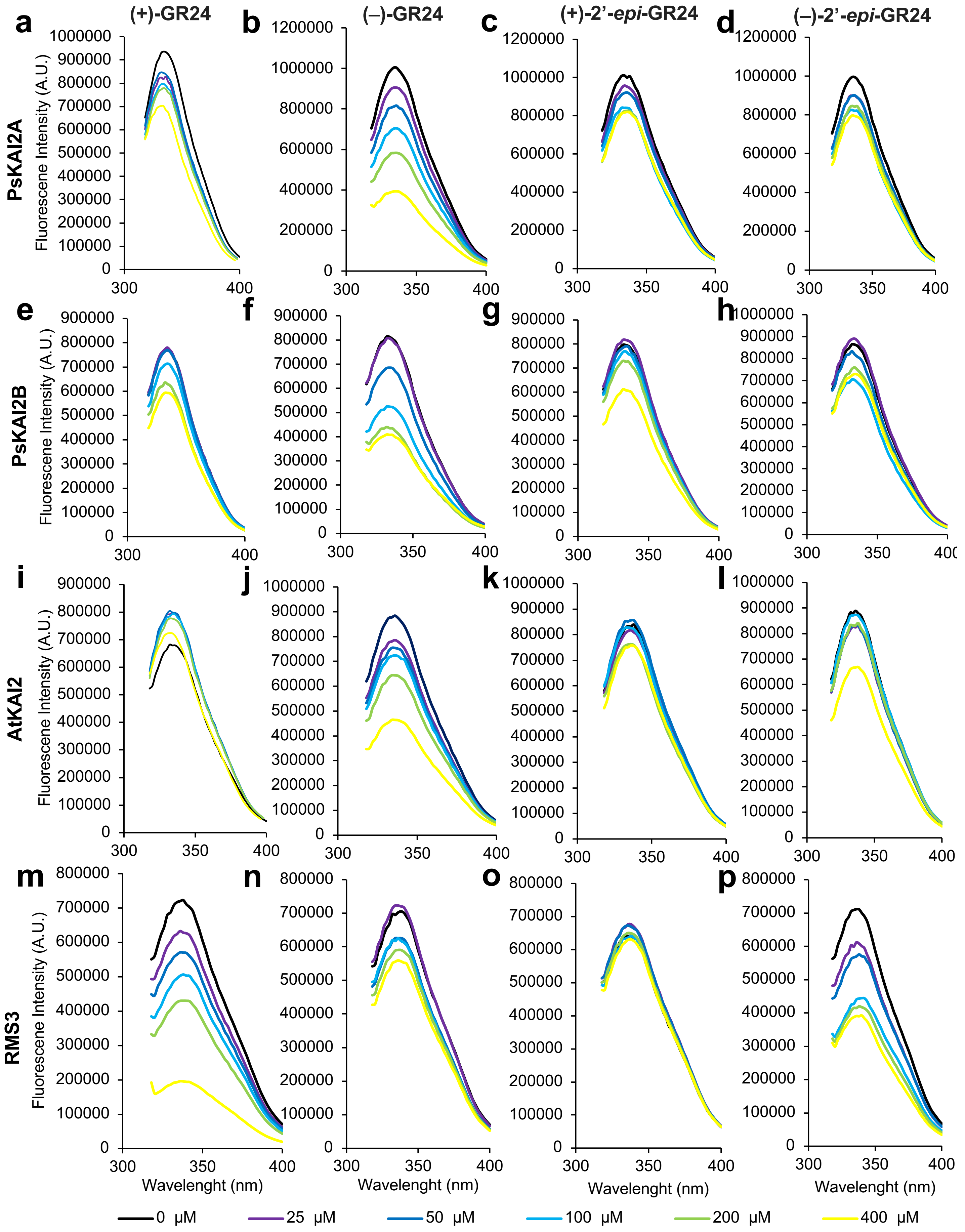

(+)-GR24
(–)-GR24
(+)-2'-epi-GR24
(–)-2'-epi-GR24
PsKAI2A
PsKAI2B
AtKAI2
RMS3
Fluorescene Intensity (A.U.)
Wavelenght (nm)
0 µM
25 µM
50 µM
100 µM
200 µM
400 µM

**Fig. S9. Intrinsic tryptophan fluorescence of PsKAI2s, AtKAI2 and RMS3 proteins in the presence of SL analogs.** Intrinsic tryptophan fluorescence of PsKAI2A (**a-d**), PsKAI2B (**e-h**), AtKAI2 **(i-l)** and RMS3 (**m-p**) proteins in the presence of SL analogs. Changes in intrinsic fluorescence emission spectra in the presence of various concentrations of (+)-GR24 (**a;e;i;m;q**), (–)-GR24 (**b;f;j;n;r**), (+)-2'-*epi*-GR24 (**c;g;k;o;s**), (–)-2'-*epi*-GR24 (**d;h;l;p;t**). Proteins (10 µM) were incubated with increasing amounts of ligand (0–400 µM, top line to bottom line, respectively). The observed relative changes in intrinsic fluorescence were plotted as a function of SL analog concentration and transformed to degree of saturation and used to determine the apparent *K*D values relevant to **Figure 2i**. The plots represent the mean of two replicates and the experiments were repeated at least three times. The analysis was performed with GraphPad Prism 8.0 Software.

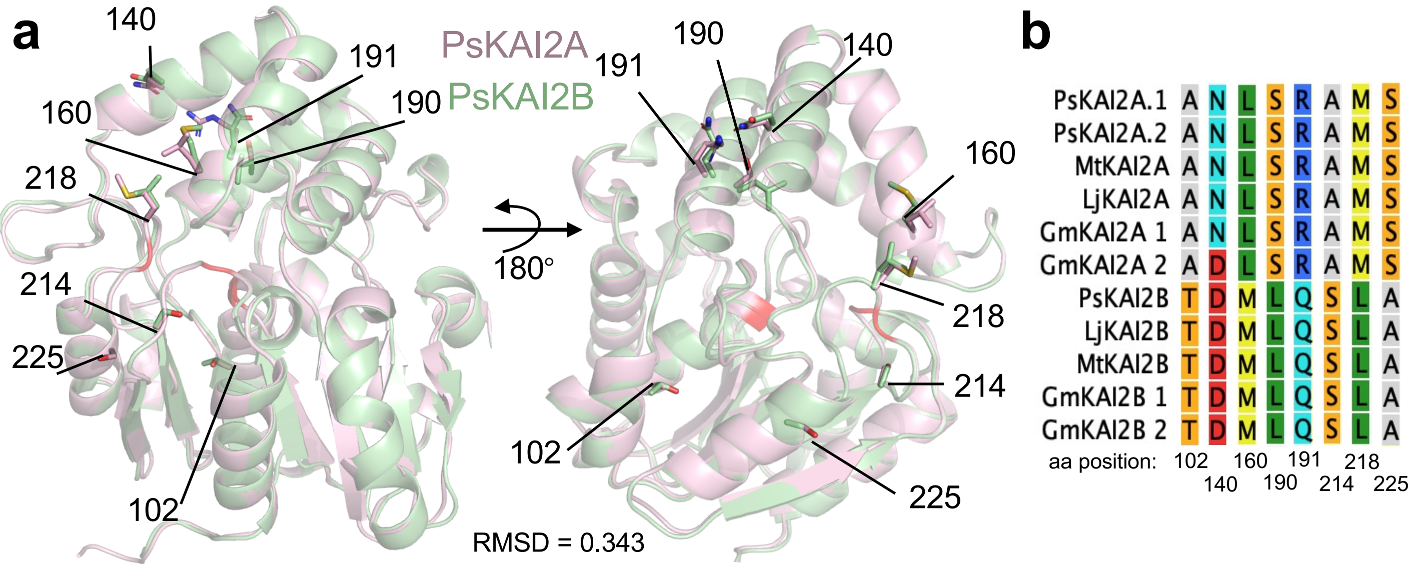

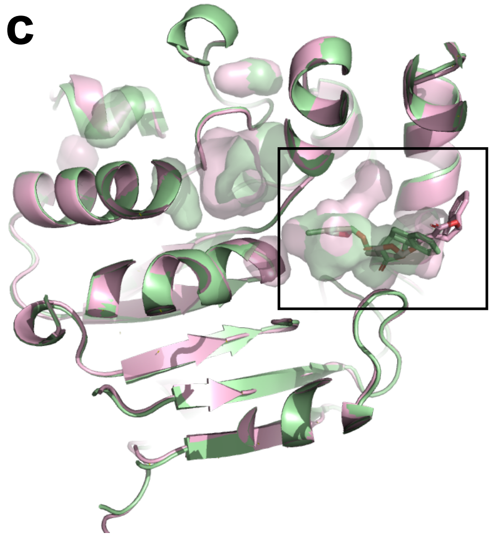

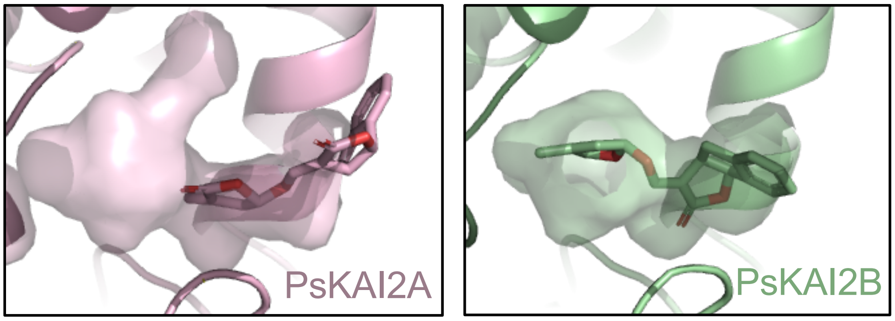

| Docking Affinity for (-)-GR24 (ΔG, kcal/mol) | |
|---|---|
| PsKAI2A | PsKAI2B |
| -4.6 | -4.9 |

**Fig. S10. Structural divergence analysis of legume KAI2A and KAI2B. (a)** Structural alignment of PsKAI2A and PsKAI2B shown in pink and light green respectively. Calculated RMSD of aligned structures is shown. Residues differentiating all legume KAI2A from KAI2B are shown on each structure as sticks and labeled with residue number. Catalytic triad is shown in red. **(b)** Residues 102, 140, 160, 190, 191, 214, 218, and 225, L190, and L218 are highlighted as divergent legume KAI2 residues, conserved among all legume KAI2A or KAI2B sequences (with the exception of D140 in GmKAI2A 2) as shown in reduced Multiple Sequence Alignment from **Fig. S1**. **(c)** *In silico* docking analysis of intact (–)-GR24 with solvent-accessible pocket shown for each structure and corresponding docking scores reported in kcal/mol.

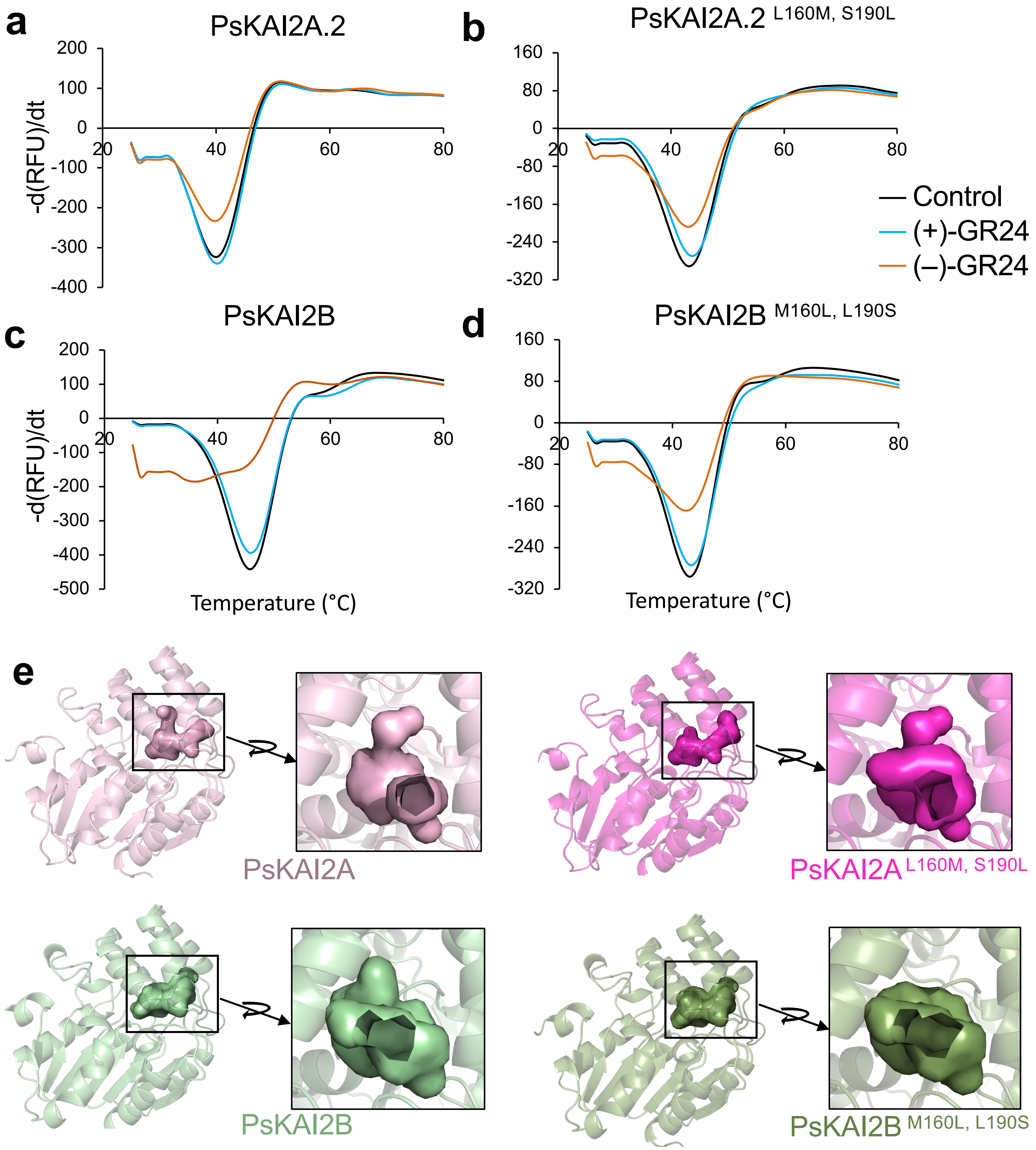

**Fig. S11. Biochemical and structural analysis of the interaction between wildtype and residue 160 and 190 swap mutant PsKAI2 proteins and (–)-GR24 by DSF.** The melting temperature curves of 10 μM PsKAI2A.2 **(a)**, PsKAI2A.2 $^{L160M,\ S190L}$ swap **(b)**, PsKAI2B **(c)**, and PsKAI2B $^{M160L,\ L190S}$ swap **(d)**, with (+)-GR24 or (–)-GR24 at the effective concentration of 62.5 μM are shown as assessed by DSF**.** Each line represents the average protein melt curve for three technical replicates and the experiment was carried out twice. **(e)** pocket surface representation of PsKAI2A model, PsKAI2B apo structure, and modelled PsKAI2A.2 $^{L160M,\ S190L}$ and PsKAI2B $^{M160L,\ L190S}$ swap mutants.

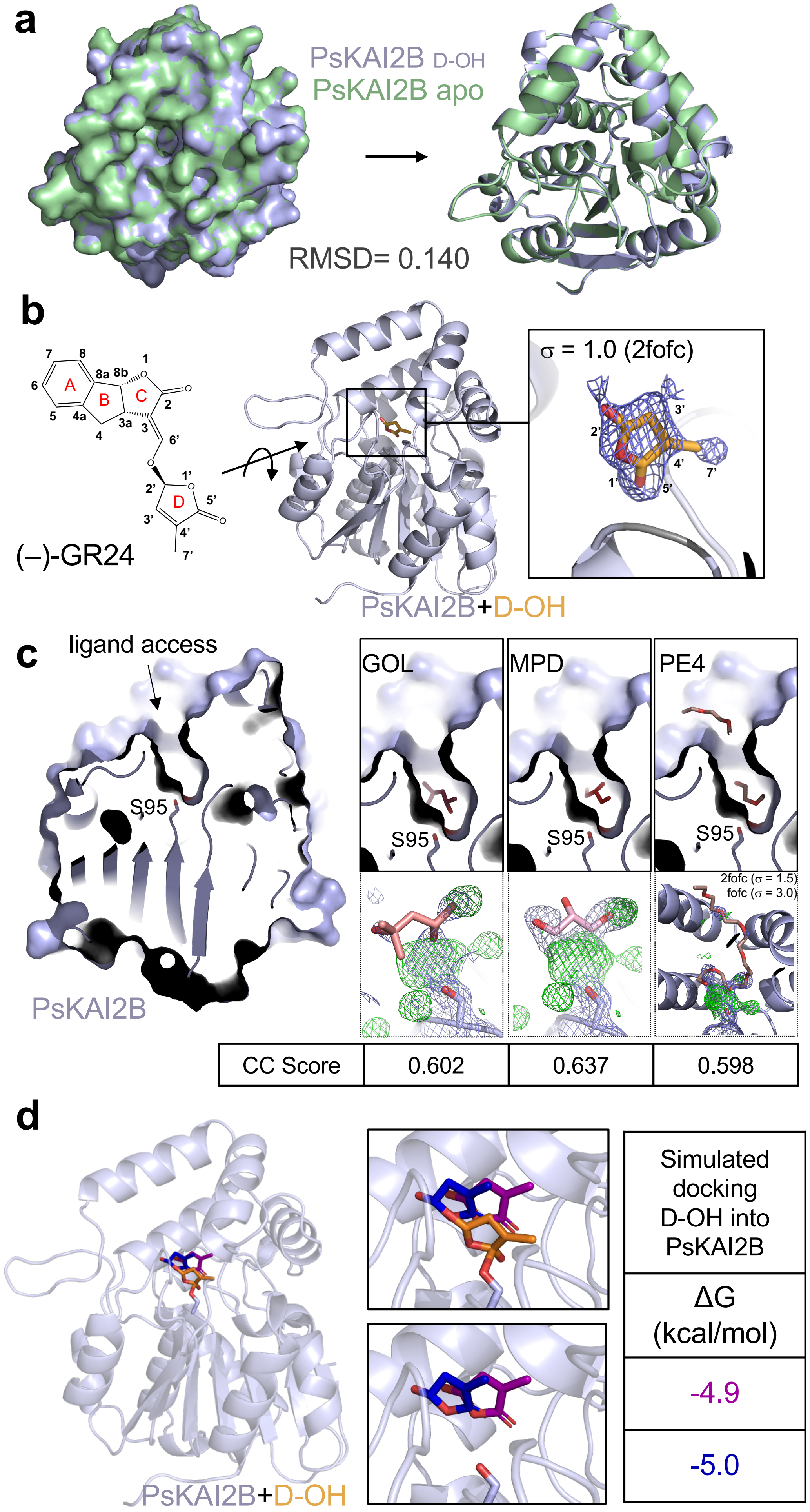
a
PsKAI2B D-OH
PsKAI2B apo
RMSD= 0.140
b
(–)-GR24
PsKAI2B+D-OH
σ = 1.0 (2fofc)
c
ligand access
S95
PsKAI2B
GOL
MPD
PE4
2fofc (σ = 1.5)
fofc (σ = 3.0)
CC Score
0.602
0.637
0.598
d
PsKAI2B+D-OH
Simulated docking D-OH into PsKAI2B
ΔG (kcal/mol)
-4.9
-5.0

**Fig. S12**

**Fig. S12. Structural interrogation of the ligand bound PsKAI2B crystal structure. (a)** Structural alignment of PsKAI2B apo (light green) and PsKAI2B in complex with (–)-GR24 D- OH (gray/blue). Calculated RMSD value is shown. Similar orientation of the superposition is shown in surface (right) and cartoon (left) representations. **(b)** Chemical structure of intact (–)- GR24 molecule with numbered carbons. Electron density mesh fit with D-OH ring of (–)-GR24. Protein structure is shown in blue/gray and ligand in orange. The electron density of the ligand is derived from 2mFoDFc (2fofc) map contoured at 1.0s. **(c)** LigandFit examination of crystallization and purification conditions reported here. Glycerol denoted: GOL, (+/-)-2-Methyl- 2,4-pentanediol denoted: MPD. Polyethylene glycol, PEG, denoted: PE4. Ligand access pocket is shown together with the ligand placed via LigandFit fitting software (top), and the obtained electron density maps following 3 cycles of Phenix refine. The electron densities shown are derived from 2mFoDFc map (2fofc, blue mesh) contoured at 1.5s and mFoDFc map (fofc, green mesh) contoured at 3s. Correlation Coefficient (CC) scores were calculated via LigandFit. **(d)** *In silico* analysis of D-OH ligand docking is shown in orange (found in the structure) and predicted orientations in navy and magenta with corresponding docking scores reported in kcal/mol.

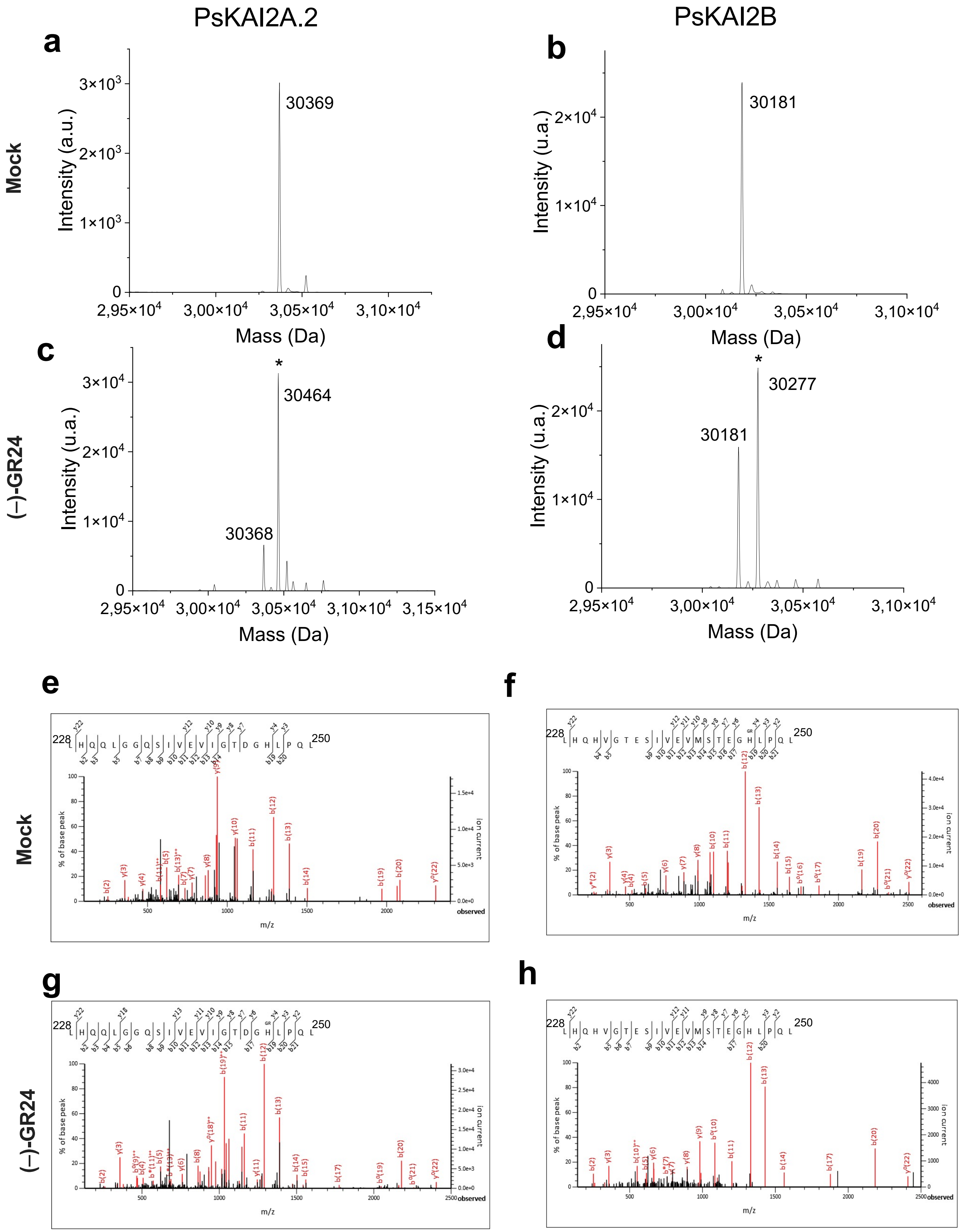

**Figure S13**. **Mass spectrometry characterization of covalent PsKAI2-ligand complexes.** A Deconvoluted electrospray mass spectra of PsKAI2A.2 and PsKAI2B before (**a-b**) and after (**c-d**). adding of ligand (–)-GR24 are shown respectively on upper and lower panels. Peaks with an asterisk correspond to PsKAI2 covalently bound to a (–)-GR24 ligand (PsKAI2-ligand). A mass increment of 96 Da is measured for two PsKAI2-ligand complexes. (**e-h**) Ligand-modified H246 amino-acids were identified by nanoLC-MSMS analyses after chymotrypsin proteolysis. Fragmentation spectra of unmodified and ligand-modified peptides are shown. Labeled peaks correspond to b and y fragments of the triple charged precursor ion. Histidine 246 residue modified by ligand is marked with GR on the sequence displayed on the top.

**Table S1. List of the mutations identified during TILLING and mutant alleles used in the study.**

| *PsKAI2A* TILLING mutants | | | | |
|---|---|---|---|---|
| **Mutant allele** | **Base position[1]** | **Protein position** | **Type of mutation** | **Protein location of the mutated amino acid** |
| *Pskai2a-1* | G266A | C89Y | Missense | |
| 2 | C333T | L111L | Silent | |
| *Pskai2a-2* | G368A | R123K | Missense | This R is conserved across KAI2s, located at the left base of the V lid in the back of a loop |
| 4 | G368A | R123K | Missense | |
| *Pskai2a-3* | G379A | D127N | Missense | This D is conserved across KAI2s, located at the left base of the V lid in the middle of a loop |
| 6 | C390T | Y130Y | Silent | |
| *Pskai2a-4* | G395A | G132E | Missense | This G is conserved across KAI2s, located on the same left loop region as *Pskai2*a-2 and *Pskai2a-3*, but are pointing more inwards so possibly closer to the entrance of the pocket |
| *Pskai2a-5* | G398A | G133E | Missense | see *Pskai2a-4* |
| 9 | G409A | E137K | Missense | |
| 10 | G412A | D138N | Missense | |
| 11 | G417A | L139L | Silent | |
| 12 | G417A | L139L | Silent | |
| 13 | G454A | A152T | Missense | |
| 14 | C465T | Y155Y | Silent | |
| 15 | G467A | G156E | Missense | |
| *Pskai2a-6* | G487A | G163R | Missense | This G is conserved across KAI2s, located at the right base of the V lid in a loop region, also pointing slightly inwards |
| 17 | G561A | L187L | Silent | |
| 18 | G676A | E226K | Missense | |
| 19 | G676A | E226K | Missense | |
| 20 | G715A | E239K | Missense | |

| *PsKAI2B* TILLING mutants | | | | |
|---|---|---|---|---|
| **Mutant allele** | **Base position[1]** | **Protein position** | **Type of mutation** | **Protein location of the mutated amino acid** |
| 1 | Before ATG | - | | |
| 2 | 44 | T15I | Missense | |
| *Pskai2b-1* | 34 | V12I | Missense | This V is conserved across all KAI2s, located on the back of the protein pointing outwards not near lid or known binding interfaces |
| *Pskai2b-2* | 254 | R85K | Missense | This R is not conserved, AtKAI2 has a K at this position, located on the back of the protein pointing outwards not near lid or known binding interfaces |
| 5 | 120 | L40L | Silent | |
| 6 | 121 | L41L | Silent | |
| 7 | 201 | Y67Y | Silent | |
| *Pskai2b-3* | 488 | G163E | Missense | see *Pskai2a-6* |
| 9 | 508 | A170T | Missense | |
| 10 | 655 | A219T | Missense | |
| 11 | 658 | V220I | Missense | |
| 12 | 664 | V222I | Missense | |
| 13 | 674 | A225V | Missense | |
| 14 | 540 | N180N | Silent | |
| 15 | 558 | A186A | Silent | |
| 16 | 663 | P221P | Silent | |

[1] base position from the ATG

**Table S2. Data collection, phasing and refinement statistics**

| | PsKAI2B (apo form, with glycerol) | (−)-GR24 D-OH - bound PsKAI2B |
|---|---|---|
| **Data collection** | | |
| Space group | C2 | C2 |
| Cell dimensions | | |
| *a*, *b*, *c* (Å) | 87.59, 71.14, 49.06 | 87.08, 71.82, 48.79 |
| α, β, γ (°) | 90, 117, 90 | 90, 117.3, 90 |
| Resolution (Å) | 43.47-1.61 (1.66-1.61)* | 43.36-2.00 (2.07-2.00) |
| $R_{sym}$ | 0.080 (0.589) | 0.082 (0.316) |
| $I / \sigma I$ | 31.01 (1.52) | 35.13 (4.11) |
| Completeness (%) | 99.2 (84.5) | 98.73 (87.53) |
| Redundancy | 6.4 (3.2) | 6.1 (4.5) |
| **Refinement** | | |
| Resolution (Å) | 1.61 | 2.00 |
| No. reflections | 34306 | 17837 |
| $R_{work}$ / $R_{free}$ (%) | 15.9/17.7 | 16.9/21.1 |
| No. atoms | 2395 | 2298 |
| Protein | 2110 | 2110 |
| Ligand/ion | 6 | 8 |
| Water | 279 | 180 |
| *B*-factors | | |
| Protein | 19.92 | 26.5 |
| Ligand/ion | 33.73 | 24.60 |
| Water | 32.07 | 32.22 |
| R.m.s. deviations | | |
| Bond lengths (Å) | 0.009 | 0.013 |
| Bond angles (°) | 0.88 | 1.03 |
| Ramachandran favored (%) | 98.51 | 98.88 |
| Ramachandran allowed (%) | 1.49 | 1.12 |
| Ramachandran outliers (%) | 0 | 0 |
| PDB ID | 7K2Z | 7K38 |

**Table S3. List of primers used in this study**

| GENE | Primer name | Sequence | Observation |
|---|---|---|---|
| Primers for qRT-PCR analysis | | | |
| *PsACTIN* | PsACTIN_F1 | GTGTCTGGATTGGAGGAT | Used in figure 1 |
| | PsACTIN_R1 | GGCCACGCTCATCATATT | |
| *PsTUB* | ST206-F | CAGAACAAGAACTCGTCATACT | Used in figure 2 |
| | ST207-R | AGCCTTCCTCCTGAACATA | |
| PsKAI2B<br>Psat4g083040 | PsKAI2B_F1 | GCCCTAAGCGTGTTACAAAC | Used in figure 2 |
| | PsKAI2B_R1 | ACTCGGTGCCGACGTGT | |
| | PsKAI2B_F2 | GCCCTAAGCGTGTTACAAAC | Used in figure 1 |
| | PsKAI2B_R2 | ACTCGGTGCCGACGTGT | |
| *PsKAI2A*<br>Psat2g169960 | PsKAI2A_F1 | CTTTGATAGTGTCGAGGACGA | Used in figure 2 |
| | PsKAI2A_R1 | GACCACCCAATTGTTGATGT | |
| | PsKAI2A_F2 | CTTTGATAGTGTCGAGGACGA | Used in figure 1 |
| | PsKAI2A_R2 | GACCACCCAATTGTTGATGT | |
| *PsDLK2*<br>Psat1g039520 | ST208-F | AGGCTTGTTCTTCTTGGTGC | Used in figure 2 |
| | ST209-R | ATCTGAACTTGCAAATCCTCCC | |
| Primers for TILLING | | | |
| *PsKAI2A*<br>Psat2g169960 | PsKAI2A_N1F | GGAAACACTTTGTACCACATCTCG | Nested 1 PCR |
| | PsKAI2A_N1R | TCCTTCATCTTGTGCCTTACTAGC | |
| | PsKAI2A_N2Ftag | ACGACGTTGTAAAACGACGATAACATG<br>GGTGCTGG | Nested 2 PCR<br>Specific primers (black nucleotides) have a M13 tag (red nucleotide) in 5'end. |
| | PsKAI2A_N2Ftag | TAACAATTTCACACAGGCAATGGGCCT<br>ATAATGGG | |
| PsKAI2B<br>Psat4g083040 | PsKAI2B_Fw1 | TTCCCTACACGACGCTCTTCCGATCTCA<br>GCCTACAGTTATCAACAAACC | Specific primers (black nucleotides) have a Illumina adaptator (red nucleotide) in 5'end. |
| | PsKAI2B_Rv1 | AGTTCAGACGTGTGCTCTTCCGATCTA<br>TTTTGGCAAACAAATCAGG | |
| | PsKAI2B_Fw2 | TTCCCTACACGACGCTCTTCCGATCTCG<br>ACAGTAATTACTTTGGAGG | |
| | PsKAI2B_Rv2 | AGTTCAGACGTGTGCTCTTCCGATCTTC<br>ACCGGAATAACAACATCC | |
| Primers for splicing variant identification | | | |
| *PsKAI2A*<br>Psat2g169960 | *PsKAI2A*_F100 | AACTCACACAGTCGGTGAAT | |
| | *PsKAI2A*_F898 | ACCCCACAATATATAGTTGTTG | |
| | *PsKAI2A*_R1233 | ATGCCACTTCCTTCATCTTG | |
| Primers for protein preparation and purification | | | |
| *PsKAI2A*<br>Psat2g169960 | PsKAI2A_F | aaaacctctacttccaatcgATGGGGATAGTGGAA<br>GAAG | |
| | PsKAI2A.1_R | ccacactcatcctccggTTACAAATCTGCCTCAA<br>GTTTC | |
| | PsKAI2A.2_R | ccacactcatcctccggTTACCTTATTGGCTCAA<br>TATTAAGTTG | |
| PsKAI2B<br>Psat4g083040 | PsKAI2B_F | aaaacctctacttccaatcgATGGGAATAGTGGAA<br>GAAGC | |
| | PsKAI2B_R | ccacactcatcctccggTCAAGCTACAATATCA<br>TAACGAATG | |
| Primers for generation of Arabidopsis transgenic lines | | | |
| *PsKAI2A*<br>Psat2g169960 | PsKAI2A_attb1 | GGGGACAAGTTTGTACAAAAAAGCAG<br>GCTtcATGGGGATAGTGGAAGAAGCA | |
| | PsKAI2A.1_attb2 | ggggaccactttgtacaagaaagctgggtcCAAATCTG<br>CCTCAAGTTTCA | |
| | PsKAI2A.2_attb2 | ggggaccactttgtacaagaaagctgggtcCCTTATTG<br>GCTCAATATTAA | |
| PsKAI2B<br>Psat4g083040 | PsKAI2B_attb1 | GGGGACAAGTTTGTACAAAAAAGCAG<br>GCTtcATGGGAATAGTGGAAGAAGC | |
| | PsKAI2B_attb2 | ggggaccactttgtacaagaaagctgggtcAGCTACAA<br>TATCATAACGAA | |
| AtKAI2<br>At4g37470 | AtKAI2_attb1 | ggggacaagtttgtacaaaaaagcaggcttcATGGGTG<br>TGGTAGAAGAAGC | |
| | AtKAI2_attb2 | ggggacaagtttgtacaaaaaagcaggcttcATGGGTG<br>TGGTAGAAGAAGC | |
